# Conserved folds enable immune antagonism across the tree of life

**DOI:** 10.64898/2026.07.21.739742

**Authors:** Lorenzo Profeta, Erin E. Doherty, Johanna M. Ebner, Gauri Deák Ünal, Emili A. Arasa-Verge, Logan Poehlman, Avi Shukla, Michael S. Kikugawa, Aniyah L. Barnett, Tristan X. Jordan, Wali Malik, Jason Nomburg

**Affiliations:** AITHYRA, Research Institute for Biomedical Artificial Intelligence, Austrian Academy of Sciences, Vienna 1030, Austria; University of Vienna, Doctoral School in Microbiology and Environmental Science, Vienna, Austria; Innovative Genomics Institute, University of California, Berkeley, Berkeley, CA 94720, USA; Department of Molecular and Cell Biology, University of California, Berkeley, Berkeley, CA 94720, USA; California Institute for Quantitative Biosciences (QB3), University of California, Berkeley, Berkeley, CA 94720, USA; Department of Microbiology, University of Washington, Seattle, Washington 98195; Rollins School of Public Health, Emory University; Department of Chemistry and Biochemistry, Hampton University, Hampton, VA 23669, USA; Department of Laboratory Medicine and Pathology, University of Washington, Seattle, WA, 98195, USA

## Abstract

Many components of human innate immunity are conserved in prokaryotes ^1,2^. While pathogens are known to evade host defenses ^3^, whether mechanisms of immune evasion share a similarly deep evolutionary or functional conservation across the tree of life remains largely unresolved. Here, we systematically explore this question by establishing The Viral Compendium (TVC), a database of over 350,000 proteins and 790,000 domains from eukaryotic, bacterial, and archaeal viruses. We find that protein structure alignments identify pan-viral clusters of proteins and domains, vastly increasing viral protein annotation rates compared to sequence-based methods. Domain co-association analysis revealed 1,351 combinations of domains that are conserved across archaeal, eukaryotic, and bacterial viruses, including fusion proteins that reconstitute the nuclease-ATPase core of the Mre11-Rad50 multiprotein complex involved in cellular DNA repair ^4^. Leveraging structural comparisons, we identify widely shared structural folds that mediate immune suppression: conserved phosphodiesterase folds encoded by both viral and bacterial pathogens that degrade nucleotide messengers, and double-stranded RNA binding domains employed across eukaryotic and prokaryotic viruses to suppress cellular sensing. Together, our results demonstrate that pathogen immune evasion is built upon conserved structural building blocks, revealing unified mechanisms and effectors of immune antagonism spanning all domains of life.

## Background

Although there are hundreds of millions of catalogued viral proteins, the biological function of most of these proteins is unclear. A major driver of this uncertainty is the rapid speed of viral evolution ^5^, which erodes primary sequence similarity and renders sequence-based comparisons ineffective. Protein structure prediction offers a powerful avenue to detect distant evolutionary and functional relationships, as protein structure is constrained by protein function and tends to be more conserved ^6^. While several pioneering studies have begun mapping the viral structural space, a comprehensive structure-based analysis spanning all viral kingdoms remains absent ^7–15^. In addition, these efforts have not mapped the distribution, function, and evolution of viral protein domains, leaving a major gap in our understanding of the basic unit of viral protein function. In our previous eukaryotic virus analysis, we leveraged structural alignments ^3^ to discover that two-histidine (2H) phosphodiesterases (PDEs) are employed by both animal viruses and bacterial viruses to evade cellular immunity mediated by cyclic GMP-AMP (cGAMP), constituting a conserved mechanism of viral immune evasion ^16,17^. This structural convergence raises a fundamental evolutionary question: is the broader landscape of pathogen host-evasion built upon a common toolkit of shared structural folds?

Here, we systematically addressed this possibility by establishing a comprehensive, unified database of predicted protein structures from throughout the virome, including over 17,000 species of eukaryotic, bacterial, and archaeal viruses. We clustered these viral proteins at the protein and domain level and found that protein structure methods vastly increase levels of viral protein annotation while identifying hidden links between structural features in divergent viral lineages. Specifically, we show that eukaryotic and bacterial viruses encode diverse PDE folds capable of cGAMP degradation. Strikingly, these viral PDEs share structural folds with enzymes employed by human bacterial pathogens such as *Mycobacterium tuberculosis* and *Vibrio cholerae*, revealing a shared mechanism of cGAMP regulation bridging bacterial and viral pathogens. In addition, we find that the double-stranded RNA binding domain (dsRBD) fold is highly pervasive throughout the viral proteome and is used by both eukaryotic and prokaryotic viruses to antagonize cellular immunity. We release our data as a comprehensive resource, The Viral Compendium (TVC) (tvc.apps.aithyra.at), enabling discovery and exploration of viral proteins and domains. Altogether, we find that viral immune antagonism is built upon conserved structural building blocks, revealing themes of immune antagonism relevant to humans and the entire tree of life.

### The shared universe of viral protein structures

To investigate the distribution of proteins and protein folds throughout viruses, we constructed a unified database of predicted structures from eukaryotic, bacterial, and archaeal viruses (**Fig. 1A**). First, we separately sequence clustered over 2.4 million bacterial virus (Bact-Vir) proteins and 250,000 archaeal virus (Arc-Vir) proteins and used Colabfold ^18^ to predict the structure of cluster representatives, followed by the incorporation of a database of over 67,000 predicted structures from eukaryotic viruses (Euk-Vir) ^3^. We then used a series of clustering steps with increasing sensitivity. First, we used MMseqs2 ^19^ to cluster protein sequences at 70% coverage and 20% identity, resulting in 312,953 sequence clusters (10,854 non-singleton clusters). Next, the predicted structures of sequence cluster representatives were clustered using Foldseek ^20^ 3Di alignments, further collapsing this dataset to 258,099 clusters (24,360 non-singleton clusters). Finally, representatives from non-singleton clusters were further clustered using the Foldseek-Lolalign ^21^, resulting in a final set of 251,654 clusters (17,915 non-singleton clusters) (**Extended Data Fig. 1A**). This dataset includes substantial viral diversity, with 4,512 Euk-Vir species, over 12,585 Bact-Vir species, and over 8,000 Arc-Vir genomes (**Fig. 1B**).

**Fig. 1.**
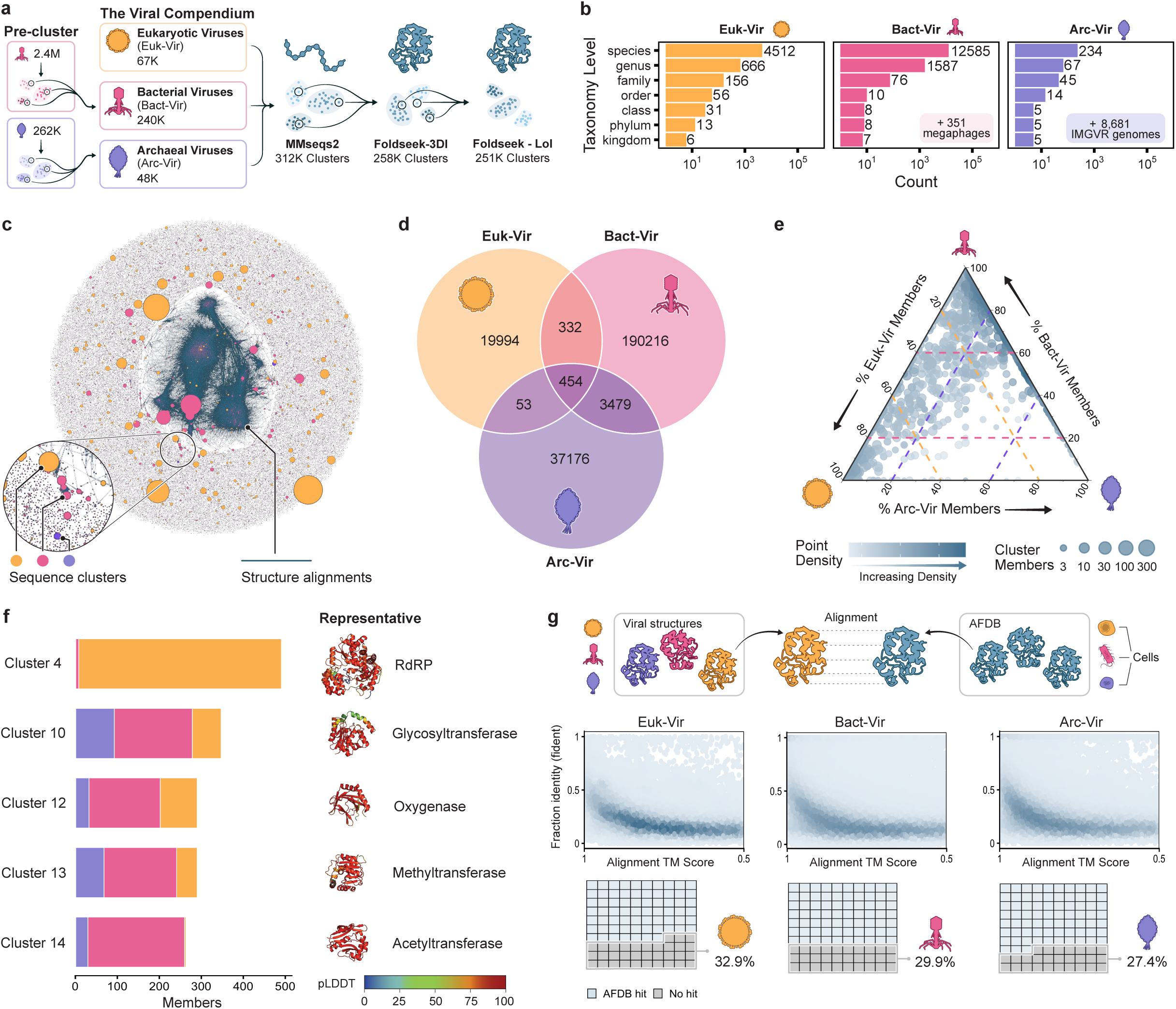
The shared universe of viral protein structures. **(A)** Cluster workflow for full viral proteins. Bact-Vir and Arc-Vir proteins were pre-clustered based on sequence similarity, and the structures of cluster representatives were predicted using Colabfold. These proteins were then jointly clustered with Euk-Vir proteins using a three-step pipeline including MMseqs2 (sequence clustering), Foldseek-3Di (structure clustering), and Foldseek-Lol (structure clustering). For this pipeline, only cluster representatives were moved to the next step. These input protein structures and all cluster information make up The Viral Compendium (TVC). **(B)** Counts of viral taxa represented in TVC. Viruses without specific taxonomic information were excluded from these counts – this includes 351 Bact-Vir megaphages, and 8,681 Arc-Vir genomes. **(C)** The network of viral proteins. Nodes represent sequence clusters, with size indicating the number of members. Edges indicate structural alignments between sequence clusters. **(D)** A Venn-diagram indicating the number of clusters that contain proteins from Euk-Vir, Bact-Vir, Arc-Vir, or combinations of virus types. **(E)** A ternary plot showing, for every cluster that contains proteins from at least two viral types, the proportion of each viral type within that cluster. Each point is a cluster, with size indicating the number of members. Position in the ternary plot indicates relative composition between Arc-Vir, Bact-Vir, and Euk-Vir derived proteins. Color indicates the density of points. **(F)** Barplots indicating the number of members in the top five clusters, by size, that contain members from Arc-Vir, Bact-Vir, and Euk-Vir. On the right, the structure of the cluster representative is indicated. **(G)** (Top) Viral structures in TVC were aligned against the AFDB. (Middle) Each point is an alignment between a viral protein and a protein in the AFDB, with Y axis indicating fraction identity and X axis indicating structural alignment TMscore. The color of the dots indicates density of points. (Bottom) These waffle plots indicate the fraction of viral proteins that do or do not have an alignment against the AFDB.

Notably, structural clustering was able to identify protein similarities at or below the limit of sequence-based homology detection ^22^ (**Extended Data Fig. 1B**), showing the benefit of protein structure comparisons for making distant connections. In this way, structural alignments can connect distinct sequence clusters, although most viral proteins fall into structurally distinct singleton clusters (**Fig. 1C**).

We found that 4,318 protein clusters contain members from multiple viral kingdoms, including 454 protein clusters with members from viruses that infect all kingdoms of life (**Fig. 1D**). The composition of these multi-kingdom clusters varies, with some containing a predominant viral kingdom and others widely represented across viral kingdoms (**Fig. 1E**). These clusters were generated from stringent clustering conditions, requiring 70% alignment coverage and not accounting for domain-level similarity between multi-domain proteins. Accordingly, we find that the largest pan-viral protein clusters of proteins above 100 residues are single-domain proteins encoding for key enzymes, including RNA-dependent RNA polymerases, glycosyltransferases, oxygenases, methyltransferases, and acetyltransferases (**Figure 1F**).

Protein structure comparisons of viral proteins in TVC with cellular folds present in the AlphaFold database (AFDB) ^23^ revealed a substantial number of alignments with robust structural similarity but very low sequence similarity (**Fig. 1G**). This is consistent with the fact that viruses have frequently acquired and adapted cellular genes over the course of their evolution ^24–26^. However, between 27.4% and 32.9% of viral proteins have no structural analog in the AFDB. These proteins tend to have a lower pLDDT (indicating lower structure prediction confidence) and shorter length than the full set of proteins (**Extended Data Fig. 1C-F**).

### Conservation of domain architectures in the viral proteome

Domains are the structural building blocks of proteins and serve as the functional units of protein evolution 27. Other efforts to study viral structural diversity across viral kingdoms focus on full-protein comparisons and clustering 8,15, limiting their ability to understand the distribution of conserved domains throughout the virome. To address this gap, we isolated protein domains directly from predicted protein structures using three domain segmentation tools 28, Chainsaw 29, Merizo 30, and UniDoc 31 (Fig. 2A). We then established a consensus, resulting in a database of 796,315 domains, of which 315,674 are well-folded and supported by at least two domain segmentation tools. Sequence clustering these domains with MMseqs2 resulted in 230,061 domain sequence clusters. Each sequence cluster representative was then moved forward for structure clustering with Foldseek, resulting in a final set of 155,648 domain structure clusters (22,445 non-singleton domain clusters). Structural searches against TED AFDB50 28 identified only 16,120 domains whose clusters had no alignments. However, most of these domains did not satisfy globularity metrics, with normalized radii of gyration above 0.365 and packing densities below 10.333, the 5th percentile of proteins included in TED-100 28,32 (Extended Data Fig. 2A, 2B). TED is based on the AFDB and is thus a summary of cellular protein domains. While there is some amount of viral contamination of AFDB and therefore TED, this result suggests that most well-folded domains in viral proteins are present in cellular proteins.

Next, we investigated the functions of these domains. We used HMMER3 hmmscan 33 to align all domains against Pfam 34, considering a sequence cluster as “sequence-bright” if at least 25% of cluster members had a Pfam alignment and “sequence-dark” otherwise. In turn, structure clusters were considered sequence-bright if at least 25% of members belong to bright sequence clusters. Brightness of members within sequence clusters is largely bimodal, with most bright clusters consisting of >90% bright members (Extended Data Fig. 2C, 2D).

Structural similarity between sequence cluster representatives revealed a dense network of sequence clusters connected by structural similarity (Fig. 2B). Notably, these structural alignments greatly outperformed sequence alignments at connecting dark and bright domains (Extended Data Fig. 2E). However, 75% of domains remained dark at the sequence level. Given this low rate of sequence-based annotation, we used Foldseek to conduct structural alignments of sequence cluster representatives against the CATH 35 and ECOD 36 databases. Here, we considered structure clusters as “structure-bright” if at least 25% of members had a CATH or ECOD alignment (Extended Data Fig. 2F, 2G). Where possible, we assigned each domain structure cluster a specific annotation label based on these alignments (Extended Data Fig. 2H, 2I). Ultimately, while only 11% of domains were sequence-bright, over 88% of domains were structure-bright (Fig. 2C). Thus, protein structure alignments provide information for a large majority of viral protein domains. We provide this information in the TVC web portal, enabling systematic assessment of viral protein function.

We found that the most abundant domain structure clusters present in all viral kingdoms include basic structural building blocks such as the jelly-roll or immunoglobulin folds, as well as enzymatic domains such as methyltransferases (Fig. 2D).

Protein function is driven by interactions between domains. Therefore, we investigated patterns of domain co-occurrence in viral proteins. For this analysis, we considered “domain groups” (Fig. 2A) as sets of domain clusters that share a structural annotation label and “protein domain architectures” as unique sets of domain groups occurring in the same protein. As a simple metric of co-occurrence, we assessed the tendency of each pair of domain groups to appear within the same protein domain architecture using odds ratios (OR), assessing statistical significance with Fisher’s exact test (Fig. 2E). This identified 26,063 significantly enriched pairs of domain groups after adjusting for multiple comparisons (Fig. 2F). Next, we investigated combinations of protein domains that are present in viruses infecting multiple kingdoms of life and that contain co-occurring domain groups. This revealed 1,351 combinations of domains present in viruses of multiple kingdoms (Fig. 2G), indicating that many domains recurrently co-occur in proteins throughout the viral kingdom of life.

One combination of domains observed in our co-occurrence analysis consists of a metallophosphatase and AAA+ ATPase domain. These domains are the main functional components of Mre11 (metallophosphatase) and Rad50 (AAA+ ATPase), the core of a conserved protein complex involved in DNA repair across the tree of life 4,37,38. To investigate further, we identified domain groups that are co-associated with the metallophosphatase or AAA+ ATPase domain groups and visualized the top 13 (Fig. 2H). This revealed a network of domain associations relating to DNA processing, including other AAA+ ATPase- and metallophosphatase-like domain groups. Mre11 and Rad50 are generally encoded by separate genes in cells as well as in viruses such as phage T437, where it is involved in host DNA degradation and recombination-dependent replication (RDR) 39,40. We identified metallophosphatase-AAA+ ATPase fusion proteins in diverse eukaryotic, bacterial, and archaeal viruses (Fig. 2I, Extended Data Fig. 3A, 3B), substantially expanding the known diversity of these fusions 41. We likewise find a variety of alternative fusion proteins. This includes Rad50-like AAA+ ATPase fusions with protein-splicing intein Hint domains and Hint-associated endonucleases, consistent with a tendency of inteins to associate with proteins involved in DNA replication and repair 42. This is curious, as these AAA+ ATPase-Hint fusions have replaced the Mre11-like exo/endonuclease 43,44 with an intein-associated endonuclease. Beyond these fusion proteins, we find that many large dsDNA viruses contain AAA+ ATPase and metallophosphatase domain groups as separate genes (Extended Data Fig. 3C), supporting the broad distribution of this complex and potential implication of these domains in RDR beyond phage T4. Together we show that domain co-occurrence patterns can yield functional and mechanistic hypotheses for poorly characterized viral proteins.

### Recurrent emergence of cGAMP PDEs in viral and bacterial pathogens

cGAS-like enzymes (cGLRs) are used by all kingdoms of life to sense and respond to viral infection, producing nucleotide messengers such as cyclic GMP-AMP (cGAMP) in response to viral cues 45–47. Eukaryotic viruses employ PDEs such as Poxin 48 and 2H-family PDEs 49–52 to degrade cGLR nucleotide signals, thus suppressing innate immunity. We recently found that 2H PDEs are employed by both animal and bacterial viruses to degrade cGAMP, constituting a shared mechanism of immune antagonism 3,16,17.

Some bacteria employ PDEs to modulate cGAMP levels of their own cGLRs or the cGLRs of their host during infection. M. tuberculosis is an intracellular bacterial pathogen of humans that can be detected by cGAS 53 and encodes the PDE CdnP to degrade 2’3’ cGAMP in infected human cells 54. V. cholerae, another human pathogen, encodes V-cGAPs that function instead by regulating 3’3’ cGAMP levels within bacterial cells, controlling bacterial chemotaxis and colonization 55–58. CdnP is a DHH-DHHA1 (DHH: Asp, His, His motif; DHHA1: DHH-associated domain 1) family PDE 59, while V-cGAPs are HD-GYP (HD: His, Asp; GYP: Gly, Tyr, Pro) family PDEs, both of which are distinct PDE folds (Fig. 3A).

Given the role of DHH and HD-GYP PDEs in bacterial infections, we investigated if viruses likewise encode DHH or HD-GYP family PDEs. Protein structure searches revealed DHH PDEs throughout eukaryotic, bacterial, and archaeal viruses (Fig. 3B). Viruses of all kingdoms likewise encode the HD PDE core, although only two hits, both from bacterial viruses, contain the GYP loop (Fig. 3B, Extended Data Fig. 4A). Next, we investigated the evolutionary relationship between viral DHH enzymes and cellular DHH enzymes such as CdnP. We used a two-step pipeline to identify cellular DHHs, using sequence search to find close cellular homologues of the viral DHH PDEs we identified, and using structure search to find diverse cellular DHH PDEs independent of sequence similarity.

Phylogenetic analysis revealed five broad groups of DHH enzymes (Fig. 3C, Extended Data Fig. 4B). Groups 1, 2, and 3 form a monophyletic clade encompassing CdnP-like enzymes from bacteria and archaea. Group 4 DHHs constitute a separate monophyletic clade notable for cellular and viral members from all kingdoms of cellular and viral life, while Group 5 is a polyphyletic clade of cellular and viral DHH PDEs. Notably, there is evidence of recurrent horizontal gene transfer of Group 1 and Group 5 DHHs from bacteria to bacterial viruses given the presence of viral proteins within well-supported cellular lineages. Likewise, Group 4 consists of a wide mixture of viral and cellular DHHs, suggestive of multiple gene-transfer events of unknown directionality. While all groups maintain the core DHH fold, there is predicted structural variation between groups (Fig. 3C). While CdnP-like PDEs have disordered residues between their DHH and DHHA lobes, Group 4 and 5 DHH PDEs maintain a distinctive ordered helix that connects the two domains. The structure of the DHHA domain likewise varies, with Group 5 DHHs being more divergent. These data show that DHH PDEs are highly diverse and spread through cellular and viral proteomes through recurrent horizontal gene transfer events.

Next, we investigated if viral DHH and HD-GYP PDEs can degrade cGAMP in the context of immune signaling. To do so, we established a cell-based reporter assay for STING signaling (Fig. 3D). Here, treatment of 293T cells with 2’3’-cGAMP, 3’3’-cGAMP, or the small molecule STING agonist diABZI 60 stimulates firefly luminescence through STING 3 (Fig. 3D). PDEs with the ability to degrade cGAMP should be able to prevent cGAMP-mediated STING activation but should be unable to prevent diABZI-mediated STING activation. The 2H PDE Acb1 can robustly degrade 3’3’- and 2’3’-cGAMP 61 and prevents cGAMP-mediated STING activation in this assay dependent on its catalytic residues (Fig. 3E). We systematically assessed the ability of viral DHH PDEs to prevent STING activation (Extended Data Fig. 4C, 4D) and found that Euk-DHH-1B, encoded by Mimivirus bradfordmassiliense (APMV), prevents STING activation by 3’3’-cGAMP dependent on its catalytic residues (Fig. 3E). Bact-Vir Ph-DHH-3B does likewise, although the catalytic mutant has non-specific pathway inhibition. Furthermore, we find that Bact-Vir Ph-HD-GYP-2 can likewise robustly prevent 3’3’-cGAMP-mediated STING activation (Fig. 3E).

To confirm that this inhibition is due to 3’3’ cGAMP degradation, we purified Euk-Vir Euk-DHH-1B and Bact-Vir Ph-HD-GYP-2 and visualized cGAMP degradation using thin-layer chromatography. We found that both PDEs can robustly degrade 3’3’-cGAMP but not 2’3’-cGAMP (Fig. 3F). In addition, we showed that Euk-DHH-1B can robustly degrade a wide variety of nucleotide messengers involved in immune-related signaling pathways (Fig. 3G), suggesting that the DHH PDE fold may be capable of broad inhibition of nucleotide signaling-based immune pathways. We found that Euk-DHH-1B (APMV gene R106) is not required for APMV replication in its amoeba host (Extended Data Fig. 5A, 5B) and its knockout does not provide a fitness disadvantage relative to wildtype (Extended Data Fig. 5C). These data suggest that Euk-DHH-1B-mediated degradation of 3’3’-cGAMP is not biologically relevant in the context of its amoeba host, consistent with the lack of known cGLRs in Acanthamoeba 62. However, Euk-DHH-1B may degrade other cyclic nucleotides during infection.

Ultimately, we find that PDEs capable of cGAMP degradation are widespread in viral and bacterial proteomes and show that shared effectors of cGAMP degradation are encoded by viral and bacterial pathogens.

### Pan-viral adaptation of the dsRBD fold for immune antagonism

Mammals employ a tiered series of sensors to detect and respond to viral dsRNA 63. These sensors use various dsRNA-binding domains, including the dsRBD fold, to bind to viral dsRNA. Eukaryotic viral proteins such as MERS NS4a and VACV E3L can inhibit mammalian dsRNA sensors, often through direct competition for dsRNA substrates 64–68 (Fig. 4A). Curiously, the dsRBD fold has likewise been adapted by phage for immune antagonism through a distinct mechanism 69. Here, the phage protein Acb4 acts as a sponge for 3’3’ cGAMP, thus repressing cGLR sensing pathways in bacteria. MERS NS4a and VACV E3L require a [KR]-X(1,3)-[KR] motif (hereafter referred to as the “KR motif”) at the N terminal side of the second alpha helix of the dsRBD fold for dsRNA binding. This motif has been lost in Acb4 (Fig. 4B).

Protein structure search revealed that dsRBDs are widespread throughout the viral proteome, with a variety of auxiliary domains (Fig. 4C). We conducted an all-by-all structural alignment of these dsRBD-containing proteins and visualized the resultant dendrogram (Fig. 4D). In addition to known eukaryotic viral dsRBDs, we found novel dsRBDs encoded by herpesviruses, iridoviruses, mimiviruses, and adenoviruses. We also found a variety of dsRBD-containing proteins in Bact-Vir. These Bact-Vir dsRBDs fall into the Acb4-like clade of single-domain proteins, apart from one phage dsRBD that contains an RNaseIII domain (Fig. 4E). Interestingly, some phage dsRBDs contain a KR motif, unlike Acb4 (Fig. 4D).

Next, we investigated if viral dsRBD-containing proteins beyond MERS NS4a and VACV E3L can inhibit dsRNA-sensing pathways. To do this, we implemented a luciferase assay that reads out MDA5-MAVS activity (Fig. 4F). In this assay, MERS NS4a and VACV E3L can suppress MDA5-driven luciferase activity dependent on their KR motifs (Fig. 4G) but fail to reduce luciferase activity downstream of MAVS overexpression, indicating that this assay is a sensitive readout of dsRNA-dependent sensing by MDA5. While the expression levels of viral dsRBD-containing proteins vary (Extended Data Fig. 6A, 6B), we found that dsRBDs encoded by herpesviruses, iridoviruses, mimiviruses, and an unclassified RNA virus can suppress MDA5 signaling while having limited ability to reduce signaling downstream of MAVS overexpression (Fig. 4G, Extended Data Fig. 6C, 6D). Mutation of each KR motif, where present, strongly reduces the ability of viral dsRBDs to inhibit the MDA5 pathway (Fig. 4G). Ph-dsRBD-10, a Bact-Vir RNaseIII, likewise can inhibit luciferase downstream of MDA5 (Fig. 4G), consistent with the reported role of Euk-Vir RNaseIII in immune antagonism in plants and amphibians 70–72 and indicating that a Bact-Vir dsRNA-binding protein is capable of cross-kingdom inhibition of eukaryotic immunity.

To confirm dsRNA binding, we purified each Euk-Vir dsRBD and determined their ability to bind a 50bp dsRNA oligo using an electrophoretic mobility shift assay. We found that many viral dsRBDs can bind this dsRNA (Fig. 4H). Mutation of the KR motif, where present, reduces dsRNA binding for most dsRBDs (Extended Data Fig. 6E).

Together, we reveal that diverse viral dsRBDs, across both eukaryotic and bacterial viruses, can suppress dsRNA-sensing pathways. More broadly, we show that conserved structural features are recurrently used for immune antagonism throughout the viral proteome, revealing a shared structural basis of immune antagonism throughout the tree of life.

## Discussion

Of the hundreds of millions of cataloged viral proteins, only a small subset has been functionally characterized. Here, we show that systematic, structure-based annotation can bypass the limitations of sequence divergence, bridge evolutionary distances and reveal hidden functional relationships across the virome. By treating protein domains as the fundamental, modular units of viral evolution, we have mapped how these structural building blocks are distributed and co-associated across the virome. This atlas, made publicly available as The Viral Compendium, provides a global resource to explore structural and functional diversity of the uncharacterized viral proteome.

A key finding of our study is that structural features capable of immune antagonism are widespread throughout the tree of life. This has several important implications. Notably, while these folds may not mediate immune antagonism in all cellular contexts, they could possess latent functional potential and be poised for immune evasion under host pressure. This is illustrated by the mimiviral DHH PDE: while dispensable for replication in its native amoebal host, it is sufficient to inhibit cGAMP signaling in human cells. Thus, these conserved structures may be especially important for rapid adaptation during viral spillover into a new host or following viral acquisition of new proteins through horizontal gene transfer. Our identification of cGAMP-degrading PDEs in viruses and pathogenic bacteria highlights a striking evolutionary convergence, demonstrating that pathogens recurrently and independently adapt a set of privileged structural scaffolds capable of immune antagonism. This indicates that the molecular rules governing host-pathogen competition are driven by uniform biochemical constraints. This concept is reinforced by the recent discovery that some bacterial pathogens of plants antagonize plant immunity using similar mechanism deployed by phages to evade bacterial immunity ^73^.

Together, our results suggest that understanding the structural features involved in immune antagonism anywhere in the tree of life can broadly inform on the pathogen-host conflict, spanning both viral and non-viral systems. Within the virome, we show that the dsRBD fold has been adapted for immune antagonism by both eukaryotic and bacterial viruses, raising the possibility that some bacterial virus dsRBDs may antagonize dsRNA-sensing within bacteria. Beyond viruses, our work demonstrates that the strategies of immune regulation and antagonism employed by pathogenic bacteria can serve as a predictive template for discovering immune-evasion mechanisms in eukaryotic and bacterial viruses.

Ultimately, our work reveals that the molecular conflict between hosts and pathogens is conducted in a highly conserved structural language. Deciphering this language through resources like TVC not only uncovers the core evolutionary principles governing host-microbe interactions, but also provides a predictive framework to anticipate, identify and neutralize emerging mechanisms of pathogenesis throughout the tree of life.

## Methods

### Preparation of protein structures

Euk-Vir proteins have been previously reported ^3^. Bact-Vir proteins: Proteins were downloaded from INPHARED ^74^ (Feb 1, 2024 release), as well as from megaphages identified by Al-Shayeb et al. ^75^. All Bact-Vir proteins were then clustered using MMseqs2 ^19^, requiring a minimum sequence identity of 20% and alignment coverage of 70%. Multi-sequence alignments (MSAs) were generated using colabfold_search against the colabfold database, downloaded May 31, 2023. Structure prediction used Colabfold ^18^, with the following settings: –stop_at_score 80, –num_recycle 3, –num_models 3,–stop_at_score_below 40, –use-gpu-relax. The model with the highest pLDDT was selected for use. Subsequently, we required that all Bact-Vir proteins downloaded from INPHARED have a bacterial host as labeled by NCBI, and discarded proteins that did not meet this requirement.

Arc-Vir proteins: All proteins available in IMGVR ^76^ were downloaded on Dec 19, 2024. We then extracted proteins with an annotated Archaeal host. In addition, we downloaded all refseq viruses with an annotated archaeal host on April 3, 2024, using NCBI Virus ^77^. These proteins were all clustered at 70% coverage and 20% identity using MMseqs2, with cluster representatives moved forward for structure prediction. MSAs were generated using colabfold_search against the colabfold database, downloaded May 31, 2023. Structure prediction used Colabfold with the following settings: –stop_at_score 70, –num_recycle 3, –num_models 5. The model with the highest pLDDT was selected for use.

### Clustering of full proteins

All Euk-Vir, Arc-Vir, and Bact-Vir proteins were clustered using MMseqs2 easy-cluster at 70% coverage and 30% sequence identity. Cluster representatives were identified, and their predicted structures were then clustered using foldseek easy-cluster using the default search mode (3Di), requiring 70% alignment coverage and alignment TMscore of 0.5. Next, representatives from non-singleton clusters were clustered using the lolalign search mode. Here, the foldseek internal clustering was not used. Instead, alignments were conducted with lolalign using foldseek easy-search, followed by filtering of the alignment file to keep alignments with a TMscore of at least 0.6 and bitscore of at least 1000. Following this filtering, the aln_cluster_greedy subcommand of SAT2 was used to conduct greedy clustering, similar in principle to the default greedy-cluster approach of foldseek cluster.

Taxonomic information for viral proteins was handled in the following way: For Euk-Vir, taxonomy IDs (taxIDs) are present in the filenames. For Bact-Vir and Arc-Vir proteins from NCBI, NCBI EntrezDirect was used to look up the taxID based on the viral genome accession. ETE3 ^78^ was then used to convert taxIDs to taxonomies. For taxIDs that resolved to a level more specific than species (often a strain), the more specific taxon was assigned the species level.

For the network presented in Fig. 1C, 50% of all sequence clusters were randomly selected and plotted.

### Alignments of full proteins against the Alphafold Database

Foldseek was used in 3Di alignment mode. As a target database, we used the Foldseek Alphafold/UniProt50 database, downloaded using the command ‘foldseek databases Alphafold/UniProt50 AF_uniprot50/db tmp’. As query, we used Foldseek databases of all Euk-Vir, Bact-Vir, and Arc-Vir structures. For visualizations of alignments in Fig. 1G, for every query we subsampled to keep the top 5 alignments by TMscore.

### Segmentation and clustering of domains

The 356,755 full protein predicted structures of TVC were fed to three different segmentation tools: Chainsaw ^29^, Merizo ^30^, and Unidoc ^31^. Merizo’s segmentation predictions were obtained with the more precise iterate method and filtered to exclude segments with less than 10 residues and N-terminal domains with less than 50 residues. The resulting domains segmented by Merizo were used as starting structures for UniDoc predictions, as it cannot classify non-domain residues.

To define the domains on which the three methods agree, we applied the consensus pipeline established by The Encyclopedia of Domains (TED, ted-v1-11-gf07e24e)^28^ with slight modifications: we corrected an off-by-one indexing error that the pipeline introduces on the consensus domains, as well as a truncation that it causes on structures with gapped residues. The pipeline assigns high, medium and low consensus tiers to the domains depending on how many methods agree on that segmentation; then, it applies a post-hoc filtering excluding segments with fewer than 6 residues and domains with less than 25 residues. This resulted in 796,315 domains, of which 467,140 had high or medium confidence and were selected for downstream analysis. These domains were further filtered to include only those with a mean pLDDT of at least 70, resulting in 315,674 final domains.

The 315,674 segmented and filtered domains were clustered using MMseqs2 easy-cluster at 70% coverage and 30% sequence identity. Cluster representatives were identified, and their predicted structures were then clustered using Foldseek easy-cluster using the default search mode (3Di), requiring 70% alignment coverage and alignment TMscore of 0.5.

### Brightness and annotation of domains

We used two different definitions of brightness: sequence-brightness was determined by sequence similarity search with HMMER3 hmmscan against the Pfam-A database; while structure-brightness was assigned based on structural similarity with respect to the CATH and ECOD databases.

Individual domains’ HMM profiles (queries) were scanned against Pfam-A profiles (release 38.1, 2025-09) with Pfam curated thresholds (--cut_ga). For each domain, query coverage against a given Pfam family was computed accounting for possible multiple non-overlapping hits on the same target, namely by taking the union of the envelope intervals of all hits to that family. Then a domain was defined as sequence-bright if it aligned to a Pfam family with 70% query coverage. Each sequence cluster was called sequence-bright if at least 25% of its members were bright. Similarly, each structure cluster was called sequence-bright if at least 25% of its substituent sequence cluster reps were bright. The structural connection between bright and dark sequence clusters was visualized through a network that was laid out with Graphviz and rendered with Cosmograph, where each cluster is a node. The edges of the network were calculated through an all-by-all Foldseek easy-search among the sequence representatives, requiring a coverage of 70% and a TMscore of 0.5.

For structure-brightness, sequence cluster representatives were searched separately against CATH (S40 non-redundant, v4.4.0, 2024-12-16) and ECOD (F40 reps, v294.2, 2026-05-05) via Foldseek search with sensitivity of 9.5, 100 maximum hits, and exact TM-align mode. The hits were then filtered with an alignment TM-score threshold of 0.5 and a query coverage threshold of 70%, and we called a sequence representative structure-bright if it had an alignment to either database. Structure clusters were defined as structure-bright if at least 25% of their substituent sequence representatives were also bright.

Next, we assigned annotation labels to a subset of bright domains and bright clusters. Specifically, individual domains were assigned the Pfam family having the highest query coverage (computed as described above), with ties broken by highest bit score, then lowest E-value. Bright sequence clusters were given a Pfam annotation label if more than half of their members had been assigned to the same Pfam family, and similarly for structure clusters by evaluating their substituent sequence representatives.

For CATH and ECOD annotations, the sequence representatives were assigned the strongest label ranking the hits by alignment TM-score, and breaking ties by query coverage, bit score, and E-value. To define the annotation label of structure clusters, we employed a threshold-walk strategy on the classification levels of the two databases independently: going from finest to coarsest classification, at each level we asked if there was a label that covered more than half of the sequence representatives, assigning it to the whole cluster if present. To aggregate multiple structure clusters into a single domain group if they shared the same label, we prioritized ECOD labels, falling back to CATH only if the cluster had no assigned label at any ECOD level.

We aligned the domains belonging to structure clusters that were sequence-dark and structure-dark against the TED AFDB50 database (https://teddymer.steineggerlab.workers.dev/foldseek/teddb_afdb50.tar.gz, 2026-02-26) with Foldseek in fast 3Di+AA mode, requiring a TM-score ≥ 0.5 and a query coverage ≥ 0.7. We retained as candidate novel domains those belonging to clusters with no confident hit. For these candidates, we computed their normalized radius of gyration and packing density using the globularity functions from CATH-AlphaFlow^79^ and compared them to the TED-100 5^th^ percentile globularity cutoffs.

### Domain co-association

Domain architectures were defined as the unique combinations of domain groups present in a full-protein sequence cluster, ignoring the number of occurrences and relative position of each domain group. Here, the same architecture can recur in multiple sequence clusters, if it is conserved.

To quantify the tendency of a domain group to co-occur on the same protein with another domain group, we used a simple pair-wise co-association test. For each observed pair of domain groups, we counted the number of architectures in which that pair was present. The enrichment of each association was calculated with the log of the Haldane-corrected odds ratio, and its significance with Fisher’s exact test, followed by BH-FDR (Benjamini-Hochberg False Discovery Rate) correction.

To identify combinations of domains that are present across multiple viral kingdoms, we analyzed all the subsets of the observed domain architectures. Specifically, we identified the combinations that were present in at least 5 different sequence clusters, whose internal pairs were all significantly enriched, and that span at least two different super-kingdoms.

To find the domain groups that were associated with the “Metallophos-AAA_23” (Mre11-Rad50) pair, we prioritized those whose association with either of the two labels (defined as poles) had the broadest support and the greatest strength. First, we identified domain groups that had a significant and positively enriched association with at least one pole, that were supported by at least two distinct full-protein sequence clusters in which they co-occurred with a significantly associated pole, and that reached a maximum log-odds ratio of at least 2. We then ranked the surviving domain groups by the product of their sequence-cluster support and maximum log-odds ratio and kept the top 13 domain groups.

To find the Mre11-Rad50-like fusion proteins and the two-gene systems, we identified all proteins containing a domain with either a metallophosphatase label or one of the labels belonging to the AAA+ ATPase constellation of domains (AAA23, AAA, Ploop, SMC, SbcC_Walker); a fusion protein falls into both sets. We then ran two structural alignments against curated references. First, the full structure of every metallophosphatase protein was aligned (Foldseek, TM-align mode) to three Mre11 references: *Pyrococcus furiosus* (PDB: 1II7), human (PDB: 9Q9H) and *E. coli* SbcD (PDB: 6S6V). A nuclease was called catalytically intact when ≥70% of the di-manganese ligand positions carried the identical reference residue (Cα within 8 Å after superposition), taking the reference with the highest fraction of conserved ligands. Second, to discriminate true Rad50 candidates we used its characteristic zinc hook, a CxxC at the coiled-coil apex ^80^. Every CxxC-bearing domain of an ATPase head protein was TM-aligned to the *Pyrococcus furiosus* Rad50 hook (PDB: 1L8D), and a hook was called confident when TM ≥ 0.55, with either the canonical CPxC motif or TM ≥ 0.70, and the parent protein was not named as an unrelated enzyme (e.g. ligase, helicase, polymerase, topoisomerase, kinase, and others). Finally, we combined the two alignments per genome. A fusion has an intact Mre11 nuclease and a Rad50 ATPase head on the same chain; a two-gene system is a genome with an intact Mre11 plus a separate Rad50 ATPase head. Candidate HINT domains were TM-aligned to the 2IN0 PDB reference (TM ≥ 0.55). We scored three splicing positions, the Block-A nucleophile (Cys/Ser/Thr), the Block-G His-Asn dyad and the C-extein +1 residue, calling a splicing-compatible intein as class 1 (all three present) or class 2 (Block-G dyad + C-extein +1 but no Block-A nucleophile) ^81^. In Fig. 2H, the Mre11 facet contains Mre11 nuclease C-terminal domain, Metallophos, Metallophos_2 and Metallo-dependent phosphatases; the Rad50 facet includes Rad50_zn_hook, SMC_N, AAA_23 and CT398 helical hairpin. In the hierarchical clustering of the matrix, the two facets are considered single entities.

**Fig. 2.**
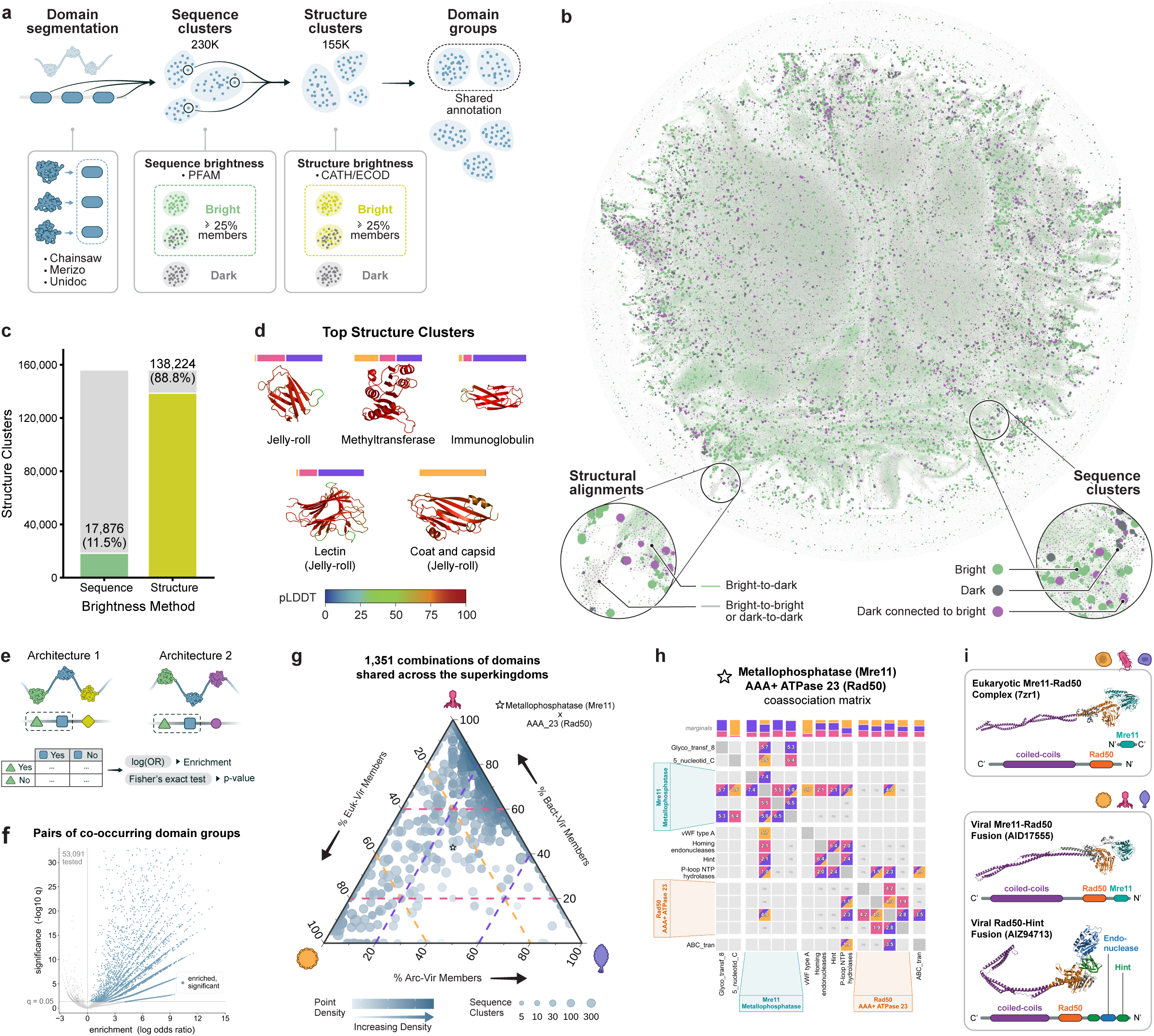
Conservation of domain architectures in the viral proteome. **(A)** Cluster workflow for viral domains. Consensus segmentation between Chainsaw, Merizo and UniDoc produced, from the full proteins, 467,140 domains supported by at least two segmentation methods, 315,674 of which had high mean pLDDT. Domain sequences were clustered with MMSeqs2, with sequence representatives further clustered with Foldseek. This resulted in 155,648 structure clusters, whose structural brightness was calculated against CATH or ECOD. The final domain groups were obtained by aggregating structure clusters that shared the same domain annotation label. **(B)** The network of viral domains. Nodes represent sequence clusters, with size indicating the number of members. Edges indicate structural alignments between sequence clusters. Bright sequence clusters against Pfam are shown in green, dark sequence clusters in grey, dark sequence clusters that connect to at least a bright one by structural similarity in purple. Edges that connect a bright with a dark cluster are green, while those that connect clusters of the same category are muted as beige on the background. For clarity, only the top 10 edges per node are displayed. **(C)** The percentage of structure clusters that are sequence-bright against Pfam is compared to the percentage of those that are structure-bright against CATH or ECOD. **(D)** The domain structures representing the 5 biggest structure clusters (in descending order, from left to right, and from top to bottom) spanning all the three superkingdoms. The colored bar indicates the proportion of sequence representatives from viruses of each type, normalized to the size of that super-kingdom. The domains are named according to the ECOD label assigned to each cluster. **(E)** Schematic representing the pair-wise co-association test of domain groups carried out over all the domain architectures in the database. (**F)** A volcano plot displaying the pairs of domain groups that have been tested for statistical enrichment with Fisher’s exact test. All the pairs that were observed co-occurring in at least one domain architecture were tested. The significant (BH-FDR corrected p-value < 0.05) and enriched (logOR > 0) pairs are colored. 265 pairs are not displayed because they sit above the upper y axis boundary. **(G)** A ternary plot displaying the 1,351 combinations of domain, derived from full protein architectures, where each pair is significant and enriched and the combination is present in at least 2 super-kingdoms. Combinations of domains were required to be supported by at least 5 distinct full proteins sequence clusters. Each point is a domain combination, with size indicating the number of sequence clusters where it is observed. Position in the ternary plot indicates relative composition between Arc-Vir, Bact-Vir, and Euk-Vir of the sequence cluster representatives. Color indicates the density of points. The starred combination includes the domain groups with the annotation labels “Metallophos” and “AAA_23”. **(H)** A co-association matrix showing the domain groups that mostly co-associate with the Metallophosphatase (Mre11) or AAA+ ATPase 23 (Rad50) domain groups. The other domain groups in the matrix were selected as the top 13 ranked by the product of sequence clusters supporting the association with either of the two poles and the maximum enrichment value achieved on either of the two poles. In each cell, the logOR of the pair is displayed and the size-normalized distribution of that pair among the sequence clusters of the three virus types is shown by coloring the cell. Co-occurring but not significant pairs are reported as ‘ns’ in grey cells. On the top, the normalized marginal proportions of each domain group are shown. **(I)** Comparison of a reference cryo-EM structure of the Mre11-Rad50 two-genes complex of *Chaetomium thermophilum* (PDB: 7ZR1), and its associated domains, with the predicted structures of two exemplar Mre11-Rad50-like fusion proteins found in The Viral Compendium. From top to bottom, Listeria phage LMTA-57 (UniProt: AID17555) and Lactobacillus LfeInf (UniProt: AIZ94713), both belonging to the Caudoviricetes.

### Identification of viral DHH-DHHA and HD-GYP PDEs

To identify viral DHH-DHHA PDEs, the structure of *M. tuberculosis* CdnP (Uniprot: Q7D6H2) was predicted using the Alphafold3 webserver. This structure was used as a query for foldseek easy-search in lolalign mode against TVC, requiring a bitscore of at least 9000 and a query coverage of at least 70%. Query coverage was calculated by determining the number of aligned residues in the alignment CIGAR string, using the SAT2 subcommand aln_cigar_to_cov, and dividing by the query length. For alignments shown in figure 3, all alignments with a query coverage of at least 70% were shown, regardless of bitscore.

**Fig. 3.**
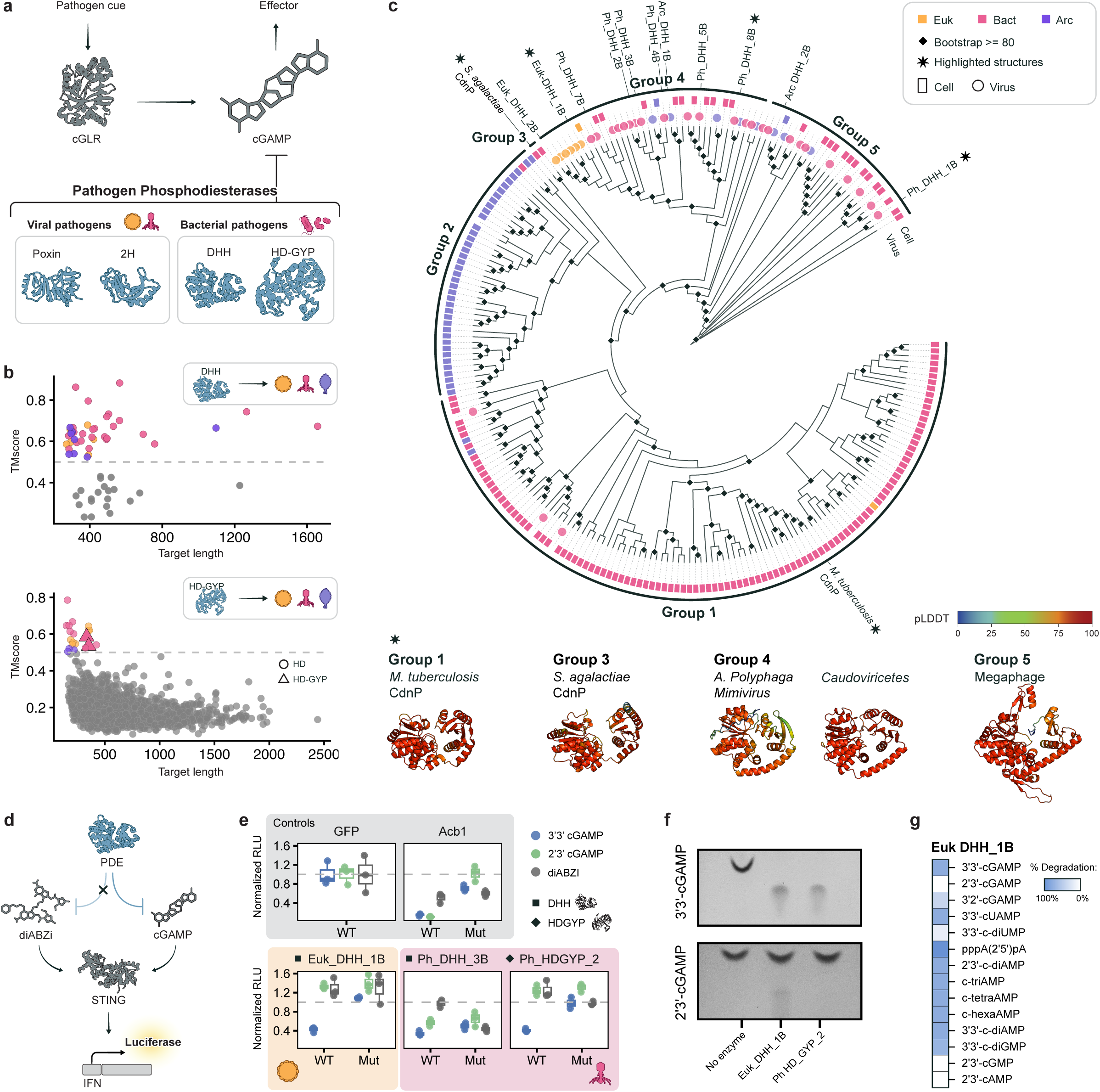
Recurrent emergence of cGAMP PDEs in viral and bacterial pathogens. **(A)** Schematic showing the targeting of cGAMP by diverse pathogen phosphodiesterases. **(B)** Protein structure alignments of *M. tuberculosis* CdnP (DHH fold, top) and *V. cholerae* V-cGAP3 (HD-GYP fold, bottom) against viral structures in TVC. Y axis indicates structural alignment TMscore, and X axis indicates the length in residues of the viral protein hit. In the bottom plot, triangles indicate hits that contain a GYP loop. **(C)** Phylogenetic tree of viral and cellular DHH PDEs. The outer ring indicates cellular enzymes, with the inner bar of circles indicating viral enzymes. Diamonds at breakpoints indicate a bootstrap value of at least 80. This tree is a cladogram, where branch length is ignored – branch lengths can be viewed in Extended Data Fig. 4B. Five structures are highlighted below – these are indicated in the tree with an asterisk. **(D)** Schematic of the cell-based cGAMP assay. Here, 293T cells are transfected with plasmids encoding a PDE, STING, firefly luciferase driven by an interferon promoter, and constitutive renilla luciferase. Cells are subsequently treated with 2’3’ cGAMP, 3’3’ cGAMP, or the small molecule diABZI. cGAMP-degrading PDEs can prevent cGAMP-but not diABZI-mediated STING activation in this system. **(E)** Cell-based cGAMP assay depicated in panel D. Y axis indicates normalized RLU, determined by the firefly/renilla ratio of each well normalized to GFP values for each STING agonist. The agonists, 3’3’-cGAMP (blue), 2’3’-cGAMP (green), and diABZI (grey) are indicated. WT indicates wildtype sequence, while Mut indicates mutation of the catalytic residues. These data are from one representative biological replicate and are a subset of the data presented in full in Extended Data Fig. 4D. **(F)** Purified Euk-DHH-1B or Ph HD-GYP-2 were incubated with 2’3’- or 3’3’-cGAMP for 16 hours, and reaction products were visualized using thin layer chromatography. **(G)** Purified Euk-DHH-1B was incubated with a variety of nucleotide messengers for one hour. Degradation of nucleotide messengers was determined through thin layer chromatography, and densitometry was conducted on a single representative replicate.

To identify viral HD-GYP PDEs, the experimental structure of *V. cholerae* V-cGAP3 (PDB: 5Z7C) was used as query for foldseek easy-search in lolalign mode against TVC, requiring a bitscore of at least 1000 and TMscore of at least 0.5.

### DHH-DHHA phylogenetic analysis

DHH PDEs were identified throughout cellular proteomes using two strategies. First, foldseek easy-search in 3Di mode was used to search the Alphafold/UniProt50 foldseek database, requiring a TMscore of at least 0.5 and query coverage of 70%. Subsequently, the structures of all resultant hits were downloaded and converted to protein sequence using the struc_to_seq subcommand of SAT2. Hit sequences were then clustered with MMseqs2, requiring coverage of at least 70% and sequence identity of at least 30%.

Second, all viral DHH-DHHA hits identified in TVC were used as query for sequence search using MMseqs2 against the Uniref50 database, requiring bidirectional coverage of at least 70% and sequence identity of at least 30%. Subsequently, the protein structures of all hits were downloaded – those without a structure in the AFDB, as well as hits of viral origin, were discarded. Structures were converted to protein sequences, and eukaryotic, bacterial, and archaeal hits were separated. Sequences were then clustered using MMseqs2 within each kingdom separately, requiring bidirectional coverage of at least 70% and sequence identity of at least 30%.

The protein sequences of all resultant sequences were collected alongside all identified viral hits, as well as supplementary sequences such as S. agalactiae CdnP (from AFDB, A0A075N0Z6). A MSA was generated using Clustal Omega ^82^ in auto mode. The resultant MSA was manually trimmed to keep the core DHH-DHHA region. IQ-Tree ^83^ was then used for phylogenetic analysis, using the -m TEST -B 1000 for model testing and bootstrapping, with the BLOSUM62+G4 ultimately being selected for use.

For horizontal gene transfer identification: Viral DHH PDEs nested within a cellular lineage were seen as evidence for cell-to-virus gene transfer. Lineages containing diverse cellular and viral DHH PDEs were seen as evidence of horizontal gene transfer of unknown directionality.

### Identification of viral dsRBDs

The TVC structure of MERS NS4a (YP_009047206) and the experimental structure of Acb4 (PDB: 9E4W, chain A) were used as queries using Foldseek easy-search in lolalign mode. Alignments required a TMscore of at least 0.6 and bitscore of at least 4500.

The dendrogram present in Fig. 4D was generated by DALI ^84^ in matrix mode, using vpSAT2 dali_matrix.sh. Here, the full protein structures of all dsRBD-containing proteins were used.

**Fig. 4.**
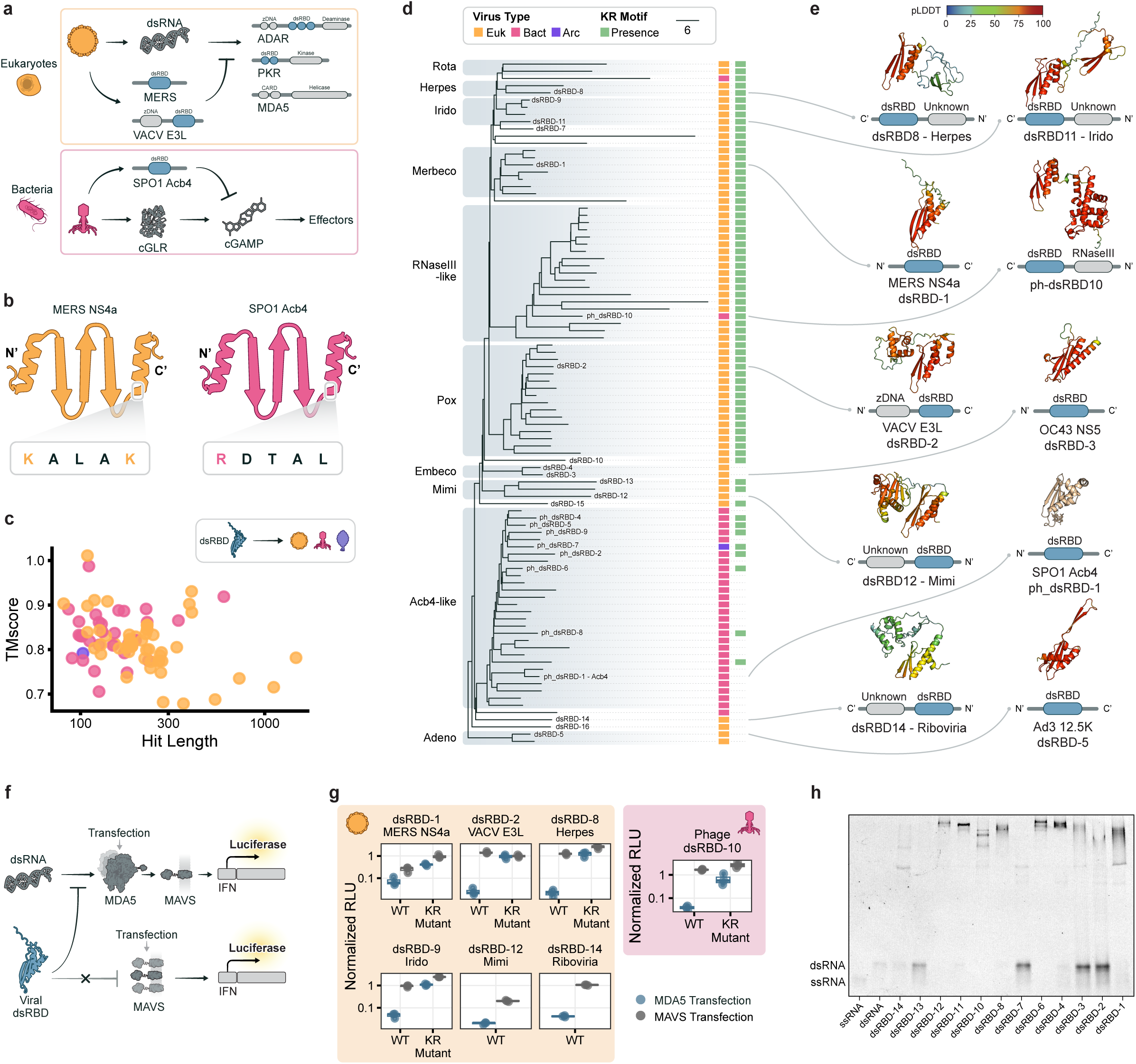
Pan-viral adaptation of the dsRBD fold for immune antagonism. **(A)** Schematic indicating the role of dsRBDs in immune antagonism and immune sensing in eukaryotes and bacteria. **(B)** Secondary structure topology of dsRBDs. The KR motif is indicated and is present in MERS NS4a but absent in SPO1 Acb4. **(C)** Protein structure alignment of MERS NS4a and SPO1 Acb4 against TVC. Y axis indicates structural alignment TMscore, and X axis indicates the length in residues of the viral protein hit. **(D)** A guide tree based on structural alignments, produced by all-by-all structural alignments with DALI. The scalebar indicates a difference of 6 in Z-score. Major clades are highlighted. Node labels indicate the dsRBDs that were experimentally tested here. Viral type and the presence of the KR motif are indicated. **(E)** Predicted structures and domain architectures of key viral dsRBDs (not to scale). **(F)** Schematic indicating the cell-based assay of MDA5 activation. Here, cells were transfected with a plasmid encoding MDA5 or MAVS, along with firefly luciferase driven by an interferon promoter, and a constitutive renilla luciferase. Viral dsRBDs are able to inhibit dsRNA-dependent MDA5-driven production of firefly luciferase, but are unable to inhibit dsRNA-independent firefly luciferase induction by MAVS overexpression. **(G)** Reporter assay to measure MDA5-MAVS activation. Cells were transfected with either MDA5 (blue) or MAVS (grey), alongside reporters and viral protein of interest. Y axis indicates normalized RLU, determined by the firefly/renilla ratio of each well normalized to GFP values separately for MDA5 and MAVS conditions. WT indicates wildtype protein. KR Mutant indicates that the KR motif was mutated to alanine. Euk-Vir and Bact-Vir data are from separate experiments. These data are one representative biological replicate and are a subset of the data presented in full in Extended Data Fig. 6C,6D. **(H)** Electrophoretic mobility shift assay of dsRBD candidates in complex with a 50bp dsRNA oligo. Unbound ssRNA and dsRNA bands are indicated.

### Cell-based luciferase assays

MDA5 assay: 96 well dishes of 293T cells (ATCC-CRL-3216) were plated at 20,000 cells/well in 150uL on day one. On day two, cells were transfected with 5ng MDA5 (Addgene 216798) or 20ng MAVS (Addgene 216798), 20ng firefly luciferase driven by the interferon beta promoter (FN-Beta_pGL3, Addgene 102597), 20ng of Renilla luciferase under the SV40 promoter (pRL-SV40, Promega E2231), and 20ng of each putative dsRBD in a custom lentiviral vector (EF1-alpha promoter) using the Mirus TransITX2 transfection reagent. On day three, firefly and renilla luciferase were measured using the Promega Dual-Glo luciferase assay system. Three wells were transfected per condition, and experiments are representative of at least two independent experiments. For normalization, the firefly:renilla ratio was determined for each well. These values were then divided by the average firefly:renilla ratio of GFP-transfected cells transfected with the same activator (MDA5 or MAVS).

cGAMP-STING assay: 293Ts were plated in 96-well plates at 20,000 cells/well in 150uL of DMEM on day one. On day two, each well was transfected with 15ng STING (pMSCV-hygro-STING R232, Addgene 102608), 20ng firefly luciferase driven by an *IFNB* promoter, 5ng renilla luciferase, and 20ng of each transgene in the pEF vector (based on Addgene 11154). After four hours, each well received a STING agonist: 0.1uM diABZI (Invivogen tlrl-diabzi-2), 50uM 3’3’ cGAMP (Biolog 117-01), 25uM 2’3’ cGAMP (Biolog 161-01). Each agonist was added directly to the cell supernatant. On day three, firefly and renilla luciferase were measured using the Promega Dual-Glo luciferase assay system. Three wells were transfected per condition, and experiments are representative of at least two independent experiments. For normalization, the firefly:renilla ratio was determined for each well. These values were then divided by the average firefly:renilla ratio of GFP-transfected cells treated with the same agonist. All data are representative of at least two independent experiments.

Detection of transgene expression: Each transgene (PDE or dsRBD) contains a HiBiT tag. On day 1, 293Ts were plated in 96 well plates at 20,000 cells per well. On day two, they were transfected with 40ng of transgene per well for PDEs, and 20ng of transgene per wel for the dsRBDs, with three wells per transgene. The next day, HiBiT levels were measured using the Promega Nano-Glo HiBiT Lytic Detection System. Average background luminescence of three wells of untransfected cells was subtracted from all values. Three wells were transfected per transgene, and experiments are representative of at least two independent experiments. All data are representative of at least two independent experiments.

The 293Ts used in this work tested negative for mycoplasma.

Working protein labels, and their corresponding protein accessions, are present in **Supplementary Table 1.**

### Protein expression and purification

Proteins were expressed and purified previously described with the following modifications ^3,16^. In brief: Expression sequences for candidate PDEs and dsRBDs were cloned into a custom pET-based vector by Gibson assembly to yield an N-terminal His10-MBP-TEV construct.

Proteins were expressed in *E. coli* Rosetta 2 (DE3) pLysS by growing cells to an OD600 of 0.4-0.6 in 2xYT (2x yeast extract tryptone) medium at 37°C, and induced with 0.5 mM IPTG (Isopropyl β-D-1-thiogalactopyranoside) after cooling to 4°C. After induction, cells expressing each protein were grown overnight at 16°C. Cells were collected by centrifugation for 20 min at 4,000 rpm, 4°C and resuspended in 20 mM Tris-HCl, pH 8.0, 10 mM imidazole, 2 mM MgCl2, 500 mM KCl, 10% (v/v) glycerol, 0.5 mM Tris (2-carboxyethyl) phosphine, and Roche protease inhibitor.

Cells were lysed by sonication, and cell lysate was clarified by centrifugation at 17,000 x g, 4 °C for 0.5 h. The supernatant was bound to Nickel-NTA affinity resin pre-equilibrated with wash buffer (20 mM Tris-HCl, pH 8.0, 500 mM KCl, 30 mM imidazole, 10% (v/v) glycerol, and 0.5 mM Tris(2-carboxyethyl) phosphine) for at least 1 h at 4°C. Supernatant was discarded and resin was washed 5 x 30 mL wash buffer (20 mM Tris-HCl, pH 8.0, 500 mM KCl, 30 mM imidazole, 10% (v/v) glycerol, and 0.5 mM Tris(2-carboxyethyl) phosphine). Protein was eluted in 5 mL elution buffer (20 mM Tris-HCl, pH 8.0, 500 mM KCl, 300 mM imidazole, 10% (v/v) glycerol, and 0.5 mM Tris(2-carboxyethyl) phosphine). Proteins were then concentrated and buffer exchanged into 20 mM Tris-HCl, pH 8.0, 500 mM KCl, 30 mM imidazole, 10% (v/v) glycerol, and 0.5 mM Tris(2-carboxyethyl) phosphine), snap-frozen and stored at −80°C.

### Analysis of recombinant protein

Purified protein was analyzed by SDS-PAGE. Samples were prepared in 1X Protein Loading Dye (50 mM Tris-HCl, pH 6.8, 15 mM EDTA, 6% (v/v) glycerol, 10% SDS, and bromophenol blue), heated at 95 °C for 3 min, loaded onto a 12% Mini-Protean TGX Precast Protein Gel (Bio-Rad) and run at 125 V until the dye front reached the bottom of the gel. Gels were stained in 30% ethanol, 10% glacial acetic acid in water, 0.1% (w/v) R-250 Coomassie and destained in 40% ethanol, 10% glacial acetic acid in water. Protein concentrations were determined by densitometric analysis of polyacrylamide gels using a bovine serum albumin (BSA) standard curve following SYPRO Orange staining.

### Recombinant phosphodiesterase activity assay

Recombinant enzymes were assayed in vitro phosphodiesterase activity against an array of oligonucleotides. All compounds were purchased from Biolog Life Science Institute or Enzo Life Sciences and used without further purification. Reactions were assayed as previously described, with the following modifications ^3,16^. Briefly, reactions were initiated by adding recombinant enzyme (17.5 µM final concentration), or an equivalent volume of reaction buffer for no-enzyme controls, to 1.25 mM oligonucleotide in reaction buffer (50 mM Tris-HCl, pH 8.0, 10 mM MgCl₂, and 100 mM NaCl).The reaction mixture was incubated at 37°C for 1 h or 16 h and stopped by heating to 95°C for 3 min. Reactions were performed in technical duplicate.

### Thin layer chromatography (TLC)

Silica gel TLC plates L × W 5 cm × 10 cm with fluorescent indicator 254 nm were spotted with 2 μL in vitro enzymatic reaction. Separation was performed in an eluent of n-propanol/ammonium hydroxide/ water (11:7:2 v/v/v). The plate was allowed to dry fully and visualized with a short-wave ultraviolet light source at 254 nm. The retention factor was calculated as the distance traveled by the observed spot divided by the distance traveled by the solvent front.

### Quantification of substrate degradation by TLC

The retention factor of non-degraded substrate bands was defined by comparison to a chemical standard. The ratio of substrate to degradation products for each timepoint was determined using densitometry of TLC images in Gel Analyzer 19.0 using a rolling ball background detection parameter with a 30% peak width tolerance. Percent degradation was determined by dividing the intensity of the substrate band by the intensity of the background for each lane. Values were normalized with the negative control. For simplicity, percent degradation values represented in the heatmap were binned such that all values ≤ 5 were assigned to bin 0, and subsequent bins correspond to increments of 10 (e.g., > 5–15 = bin 1, > 15–25 = bin 2, > 25–35 = bin 3, etc.).

### Preparation of RNA substrates for binding analysis

Cy5-fluorescently labelled RNA (ED_o_222) was purchased with PAGE gel purification from Gene Link Inc. Reverse complementary RNA (ED_o_223) was purchased from Integrated DNA Technologies, purified by denaturing polyacrylamide gel electrophoresis and visualized by UV shadowing. The band was excised from the gel, crushed and soaked overnight at 4°C in 0.5M NaOAc, 0.1% sodium dodecyl sulfate (SDS), and 0.1 mM EDTA. Polyacrylamide fragments were removed with a 0.2 μm filter, and the RNA was precipitated from a solution of 75% EtOH at −70°C for 4 h. The solution was centrifuged at 13,000 rpm for 20 min and supernatant was removed. The resultant RNA pellet was lyophilized to dryness, resuspended in nuclease-free water, and quantified by absorbance at 260 nm. Duplex RNA was hybridized at 100 nM in 1X TE and 100mM NaCl at a 1:1.2 ratio by heating to 95°C for 5 min and slowly cooling to room temperature. RNA duplex was stored at 4°C until use, or re-hybridized if frozen.

### Nucleotide sequences

Cy5-fluorescently labelled RNA; ED_o_222: 5’-Cy5-GUAUCAUCGUCAAAACCAAUUAACCAAUAUCACAUUAACUCGCAGCGUGC-3’

Complementary RNA; ED_o_223: 5’-GCACGCUGCGAGUUAAUGUGAUAUUGGUUAAUUGGUUUUGACGAUGAUAC-3’

### dsRNA binding analysis of candidate dsRBDs

Freshly hybridized dsRNA (5 fmol) was combined with 1 µg purified dsRBD candidates at 30°C for 30 min in 20 mM Tris-HCl, pH 8.0, 500 mM KCl, 30 mM imidazole, 10% (v/v) glycerol. To the reaction mixture was added 50% (v/v) dyeless formamide loading buffer (80% formamide in 1× Tris-borate EDTA (TBE) and 1mM EDTA). The resultant mixture was loaded into a 12% Mini-Protean TGX Precast Protein Gel (Bio-Rad) at 4°C and run at 150V for 90 minutes in native gel running buffer (25 mM Tris-Cl, pH 8.8, 250 mM glycine). Gels were imaged using an Amersham Typhoon Biomolecular Imager (Cytiva).

### Cells and viruses for mimivirus work

*Acanthamoeba polyphaga* (ATCC 30461) and *Acanthamoeba castellanii^Neff^* (ATCC 30010) were obtained from ATCC and maintained at 27°C in PYG (20g L^−1^ peptone, 2g L^−1^ yeast extract, 9g L^−1^ glucose, 4mM MgSO_4_, 0.4mM CaCl, 50μM Fe(NH_4_)_2_(SO_4_)_2_, 2.5mM Na_2_HPO_4_, 2.5mM KH_2_PO_4_, 4mM sodium citrate) supplemented with 1% (v/v) penicillin-streptomycin (Gibco, 15140122). Wildtype *Mimivirus bradfordmassiliense* (APMV; Chantal Abergel, IGS Marseille) and ΔR106 APMV were purified using CsCl ultracentrifugation method as demonstrated in ^85^. Briefly, a confluent layer of *A. castellanii* (∼3 × 10^^7^ cells) was infected with virus at an MOI of 1 and incubated at 27°C for complete lysis. The lysates were centrifuged at 500xg for 10 min to pellet down debris and the supernatant was centrifuged at 16000xg for 20 min to pellet down the virus particles. The particles were washed once with PBS and then resuspended in 1 mL of PBS. This was overlaid onto a CsCl gradient (made of 1.5 g/cm^3^ > 1.4 g/cm^3^ > 1.3 g/cm^3^ > 1.2 g/cm^3^) and centrifuged at 30,000 RPM for 2 h in a SW-41Ti swinging bucket rotor. Virus particles formed a band at 1.3 g/cm^3^ region of the gradient. This band was harvested and then washed with PBS three times to remove remaining CsCl. Once purified the viruses were resuspended in PBS. All viral titers were calculated by TCID_50_ method.

### Knock-out of R106 by homologous recombination

R106 was knocked out by using the protocol from ^86^. Briefly, a recombinant plasmid was constructed by sequential cloning of 5’ and 3’ flanking sequences of gene R106 as homologous arms, in the vector vHB47 with NAT (Nourseothricin Acetyltransferase) cassette. The knockout plasmid was linearized by digestion with HindIII and NotI. 4µg of linearized plasmid was transfected into *Acanthamoeba* using Polyfect (Qiagen), and infected with APMV (MOI 0.1) 1 hr after transfection. After cell lysis, viral progeny was screened for recombinants by diagnostic PCR (TerraPCR, Takara). Recombinant progeny were passaged under Nourseothricin selection to eliminate the wild-type viruses for two passages, and this was followed by amplification of recombinants without selection. After amplification, recombinants were isolated by clonal selection in 96-well format. Purity of the final ΔR106 isolate was validated by diagnostic PCR.

### Mimivirus competition assay

2 x 10^6^ *A. castellanii* were seeded in 10 cm dishes (n = 3) with 10 mL of PYG and infected with 1:1 mix of WT and ΔR106 APMV for a combined MOI of 1 (MOI 0.5 of each). One hour after infection, *A. castellanii* were washed with PYG to get rid of uninternalized virus and fresh PYG was added. Infection proceeded for 48h after which the cell debris was pelleted (5 min, 500xg) and clarified supernatants harvested. These supernatants constituted passage 1 (P1). 100 μl of P1 supernatant was used to infect a fresh confluent layer (2 x 10^6^ cells) of *A. castellani* while the rest of it was used to extract genomic DNA for qPCR analysis. This was continued until passage 5. To quantify the amount of each virus in the population three sets of qPCR primers were used. One to specifically amplify the ΔR106 mutant (NAT primers); another to specifically amplify WT APMV (R106 primers), and a final one to measure the total amount of APMV (L250, a gene shared by both). qPCR was used to measure genome copy number based off a standard curve of genomic DNA isolated from purified WT or ΔR106 virions. For each passage, the amount of WT (R106) or ΔR106 (NAT) genome copies was normalized to the total number of genome copies (L250) in that passage. Normalized values are plotted for each passage. Each biological replicate (n = 2) was performed in technical triplicate. Biological replicates are plotted.

Primers used in this study:

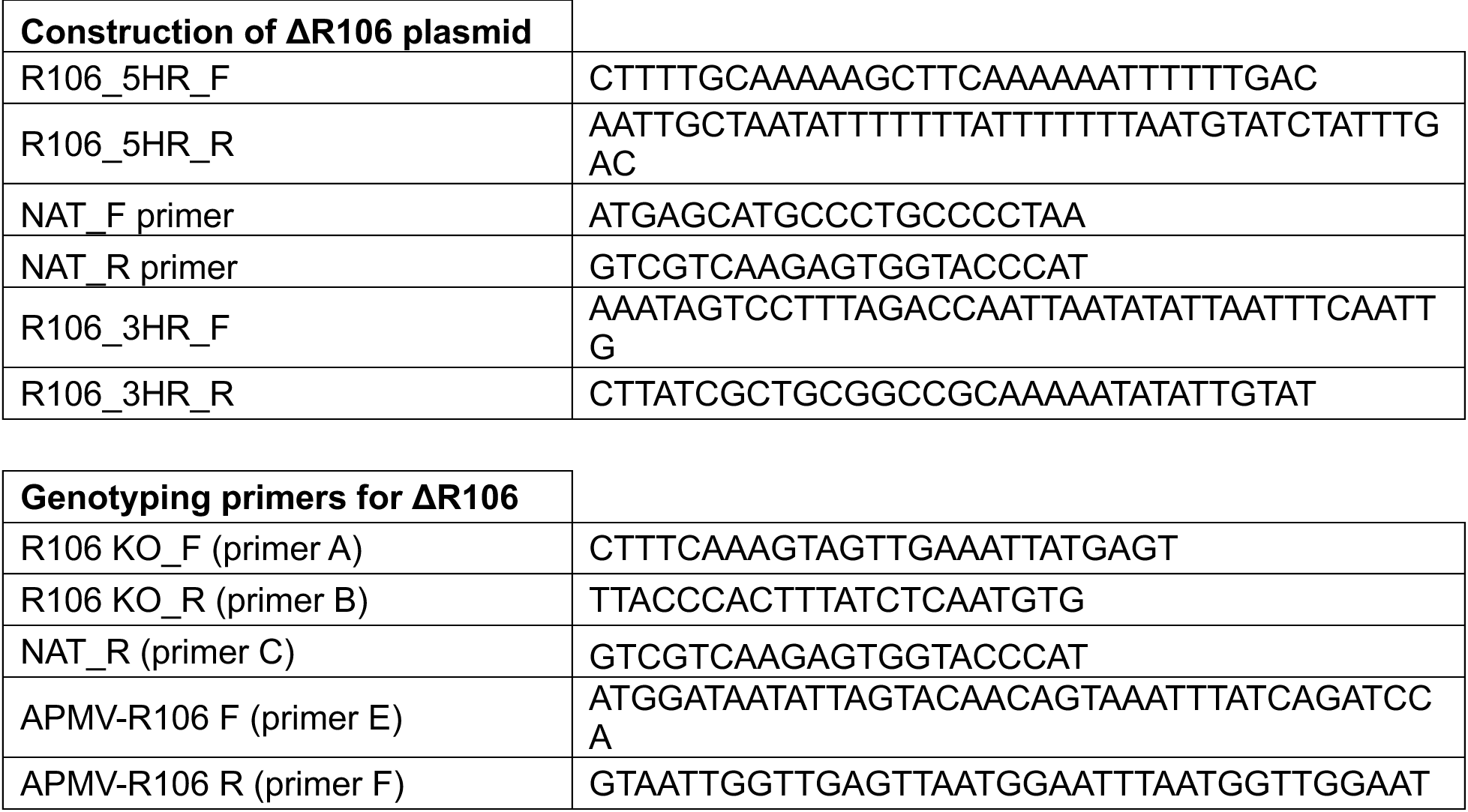
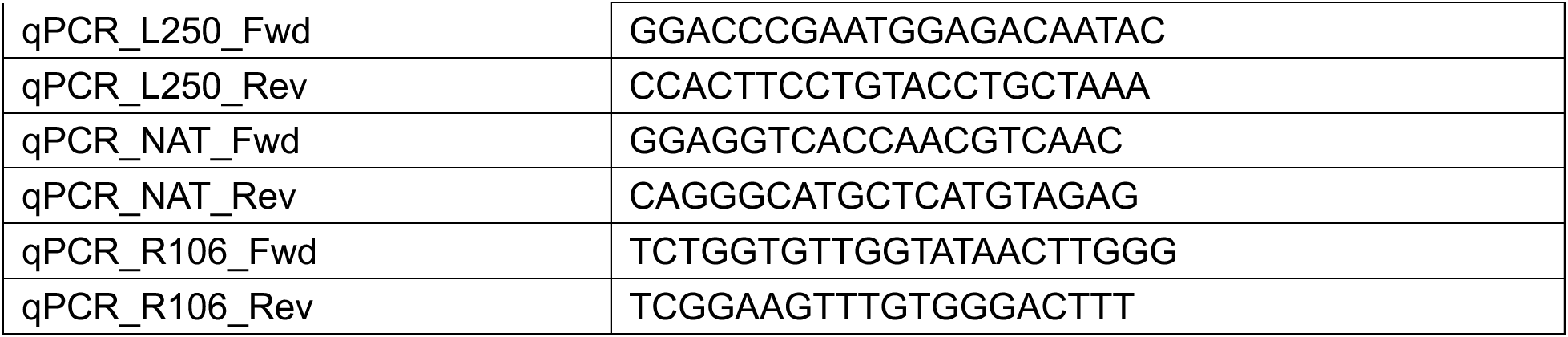

### Data analysis and plotting

Plotting for main figures 1, 3, 4 and Extended Data Figures 1, 4, and 6 were conducted in R version 4.5.2, with the exception of Fig. 3G which was assembled in GraphPad Prism. Plotting for most panels of main figure 2 and Extended Data Fig. 3 were conducted in Python version 3.11.15, except for Fig. 2C and 2G which were plotted in R. Figures were assembled in Illustrator.

### Software versions

Unless otherwise defined, software versions are as follows:

- SAT2: released here
- vpSAT2: released here
- MMseqs2: 18.8cc5c
- Colabfold: 1.5.2
- Foldseek: 799792f3799d76688f51122871229a20ad859dc0 (2025-11-12 release)
- EntrezDirect: 24.0
- ETE3: 3.1.3
- Clustal Omega: 1.2.4
- IQ-Tree: 3.0.1
- DALI: DaliLite v5.0
- Chainsaw: unversioned release (git-cloned on 2024-06-12)
- Merizo: 1.0.0
- Unidoc: unversioned release (2022-04-19)
- HMMER: 3.4
- Scipy: 1.17.1
- Statsmodels: 0.14.6
- Numpy: 2.4.6
- Pandas: 3.0.3
- Graphviz: 14.1.2
- Matplotlib: 3.10.9
- Seaborn: 0.13.2

## Data availability

Viral protein structures, domains, and clusters can be interactively explored in The Viral Compendium, available at tvc.apps.aithyra.at. This includes options to bulk download all data. TVC supports protein structure and sequence searches against TVC structures and supports plain-language queries about all protein structures and domains, informed by alignments against Pfam, CATH, and ECOD. TVC also provides a Model Context Protocol (MCP), enabling programmatic access for agents and notebooks.

Protein structures in TVC are also available for bulk download via Zenodo: https://doi.org/10.5281/zenodo.10291580 (Previously released Euk-Vir full protein structures), https://doi.org/10.5281/zenodo.21429812 (Bact-Vir and Arc-Vir full protein structures, and all protein domains from Euk-Vir, Bact-Vir, and Arc-Vir).

## Code availability

Code required to reproduce all analyses and figures are available at https://github.com/NomburgLab/2026_Profeta_et_al. This repository includes comprehensive workflow descriptions and pipelines that explain most processing steps and point to necessary intermediate files (available on Zenodo).

SAT2 is available here: https://github.com/NomburgLab/SAT2. vpSAT2 is available here: https://github.com/NomburgLab/vpSAT2.

Backups of SAT2 and vpSAT2, as well as all plotting and intermediate data needed to reproduce this work, are available on Zenodo: https://doi.org/10.5281/zenodo.21457174.

## Author contributions

Conceptualization: J.N.

Methodology: L.P., J.N.

Software: L.P., J.N.

Formal Analysis: L.P., E.E.D., A.S., J.N.

Investigation: L.P., E.E.D., J.M.E., G.D.U., E.A.A.-V., L.P., A.S., M.S.K., A.L.B., W.M., J.N.

Writing – Original Draft: L.P., J.N.

Writing – Review and Editing: L.P., E.E.D., G.D.U., J.N.

Supervision: T.X.J., W.M., J.N.

Funding acquisition: J.A.D, T.X.J., J.N.

## Acknowledgements

The authors thank Joana Carvalho for her assistance with figure and schematic design, and Stephan Stadlbauer for the development of the TVC portal. We also thank Jennifer Doudna for helpful discussions and provision of the computational resources to fold protein structures in TVC.

## Extended Data Fig. legends

**Extended Data Fig. 1 A.** Number of clusters at each clustering stage. Y axis indicates the number of clusters. The dark bar indicates the number of non-singleton clusters, while each lighter bar indicates the number of singleton clusters **B.** Distribution of structural alignments between viral proteins during 3Di clustering. Each point is an alignment between two viral proteins. The Y axis is the fraction sequence identity, and the X axis the structural alignment TMscore. The color of the dots indicate the point density. The distributions on the top and right of the plot correspond to point density along the X and Y axis, respectively. **C-F**. All plots have the density on the Y axis, separated by viral type. Panels (C) and (E) indicate the distribution of protein pLDDTs, with panels (D) and (F) indicating the distribution of protein lengths. (C) and (D) show all proteins, regardless of presence or absence of structural alignments against the AFDB. (E) and (F) show only proteins with no AFDB alignments.

**Extended Data Fig. 2** A. The density distribution of the normalized radius of gyration of members of dark clusters that did not have any hit in TED AFDB50. The TED-100 5^th^ percentile, 0.356, is highlighted. B. The density distribution of the packing density of members of dark clusters that did not have any hit in TED AFDB50. The TED-100 5th percentile, 10.333, is highlighted. C. The distribution of the proportion of sequence-bright (Pfam-bright) domains in the 28,173 non-singleton sequence clusters. The threshold used to call a sequence cluster bright, 25% of its member domains, is shown in the plot. D. The density distribution of the proportion of sequence-bright (Pfam-bright) sequence clusters in the 22,445 non-singleton domain structure clusters. The threshold used to call a structure cluster bright, 25% of its substituting sequence clusters, is shown in the plot. E. A bar chart showing the sequence-brightness (Pfam-brightness) of individual domains. Sequence-aligned to bright represent dark domains that share their sequence cluster with a bright one; structure-aligned to bright represent dark domains belonging to dark sequence clusters that share a structure cluster with a bright sequence cluster. F. The density distribution of the proportion of CATH-bright (having a confident structural hit in CATH) sequence representatives in the 22,445 non-singleton domain structure clusters. The threshold used to call a structure cluster bright, 25% of its substituting sequence clusters, is shown in the plot. G. The density distribution of the proportion of ECOD-bright (having a confident structural hit in ECOD) sequence representatives in the 22,445 non-singleton domain structure clusters. The threshold used to call a structure cluster bright, 25% of its substituting sequence clusters, is shown in the plot. H. A schematic representing the process of assigning a PFAM family at the different clustering levels: individual domains are assigned the family of the best hit covering at least 70% of the query domain, if any; sequence clusters are assigned the family that covers more than half of them; structure clusters are assigned the family that covers more than a half of the sequence representatives. I. A schematic representing the process of independently assigning CATH and ECOD labels at the different clustering levels: sequence representatives get assigned the label of the best hit with an alignment TM-score of at least 0.5, if any; structure clusters get assigned the label at the finest classification level that reaches more than half of the sequence representatives, with the highest proportion of themA covered.

**Extended Data Fig. 3** A. A table describing the Mre11-Rad50-like fusion proteins found in The Viral Compendium, along with the intein variants of that complex. The confidence is high when the Rad50 evidence includes the presence of a Rad50 zinc hook or a domain annotated as SbcC Walker B in the protein, which are both highly specific Rad50 signals; medium confidence is assigned to proteins that only show the Rad50 AAA ATPase head. Only proteins containing a metallophosphatase with at least 6 conserved metal-coordinating residues out of the 8 present in the Mre11 references are reported; the alignment TM-score with the best reference is also shown. For the intein variant of the complex, we report the intein class, which depends on the Block-A, Block-G and C-extein +1 residues. The two classes are reported as splicing-compatible, and the alignment TM-score with the intein reference (2IN0) is also shown. Then, we show the LAGLIDADG-acid position (Asp20) check and the alignment TM-score to the reference homing endonuclease (1G9Y). Lastly, we report the presence of an intact Mre11 metallophosphatase on the same genome. B. The predicted structures of the Mre11-Rad50-like fusion proteins and their fusion variants that were not shown in main Figure 2 are displayed. The metallophosphatase domain is colored in teal, the AAA ATPase head in orange and the coiled coils in purple. For the intein variants (last row), the intein (HINT) domain is shown in green and the homing endonuclease in blue. C. A bar chart showing the number of different viruses that display a high or medium confidence Mre11-Rad50-like complex as two separate genes, a fusion chain, or either of the two core components alone on the genome. In the order of fusion proteins, two-gene complexes, and solo components, viruses that are already counted in the previous category are excluded from the next one. The bar colors are stacked following the proportion of virus types, separately highlighting the large ds-DNA viruses, specifically *Nucleocytoviricota* (NCLDVs) for eukaryotic viruses and *Caudoviricetes* for bacteriophages.

**Extended Data Fig. 4** A. Structural alignment between V-cGAP3 (grey, Uniprot: Q9KL18) and phage HD-GYP (blue, protein: NODE_12_length_310542_cov_175.846709.1 X X 00139). Both HD motifs are indicated in orange, and the GYP loops are indicated in red. B. Sequence based phylogenetic tree of DHH PDEs across cells (outer ring of squares) and viruses (inner ring of circles). Diamonds indicate bootstrap values of at least 80. Leaf labels indicate the DHH PDEs tested here. This is the same tree as in Fig. 3C, but with branch lengths considered. The scalebar indicates differences per residue. C. HiBiT expression assay for DHH PDEs tested here. Y axis indicates the hibit luminescence after subtracting the background signal from transfected cells. X axis indicates the transgene, with three wells measured per transgene. D. cGAMP-STING cell-based assay results for all DHH PDEs. Reporter assay to measure STING activation. Y axis indicates normalized RLU, determined by the firefly/renilla ratio of each well normalized to GFP values for each STING agonist. The agonists, 3’3’-cGAMP (blue), 2’3’-cGAMP (green), and diABZI (grey) are indicated. WT indicates wildtype sequence, while Mut indicated mutation of the catalytic residues. These data are from one representative biological replicate. A subset of these data is presented in Fig. 3E.

**Extended Data Fig. 5** A. Euk-DHH-1B (R106) knock-out by homologous recombination. The mixed population at p0 was passaged four times with (green) or without (purple) nourseothricin selection before clonal isolation of pure progenies. The pure clonal population was confirmed by genotyping using different set of primers (a + b = 2671bp for ΔR106 and 2293 for WT, a + c = 1068 bp for integration of ΔR106 KO cassette, e + f = 1266 bp for R106 gene in WT). B. The number infectious viruses released after 16 h of infection done at MOI of 0.01 was estimated by end-point dilution and the TCID50 was calculated by standard statistical Reed and Muench method. The experiment is done in technical and biological triplicates (n = 3). AC = *Acanthamoeba castellanii^Neff^*; AP = *Acanthamoeba polyphaga*. C. Competition assay between wild-type and ΔR106 APMV across five passages. qPCR of L250 was used to measure the total APMV DNA. qPCR of R106 or the NAT cassette was used to measure the abundance of WT or ΔR106 virus, respectively. The ratio of R106 or NAT to L250 at each passage is shown. The experiment is done in technical triplicates and biological duplicates (n = 2).

**Extended Data Fig. 6 A-B.** HiBiT expression assay for the Euk-Vir (A) and Bact-Vir (B) dsRBDs tested here. Y axis indicates the HiBiT luminescence after subtracting the background signal from transfected cells. X axis indicates the transgene, with three wells measured per transgene. **C-D.** MDA5 cell-based assay for the Euk-Vir (C) and Bact-Vir (D) dsRBDs investigated here. Cells were transfected with either MDA5 (blue) or MAVS (grey), alongside reporters and viral protein of interest. Y axis indicates normalized RLU, determined by the firefly/renilla ratio of each well normalized to GFP values separately for MDA5 and MAVS conditions. WT indicates wildtype protein. KR Mutant indicates that the KR motif was mutated to alanine. Euk-Vir and Bact-Vir data are from separate experiments. A subset of these data are presented as Fig. 4G. **E**. Electrophoretic mobility shift assay of wildtype and KR-mutant dsRBDs with dsRNA. Unbound ssRNA and dsRNA bands are indicated.

