## Extended Data Figures for "Conserved folds enable immune antagonism across the tree of life"

**Extended Data Fig. 1**

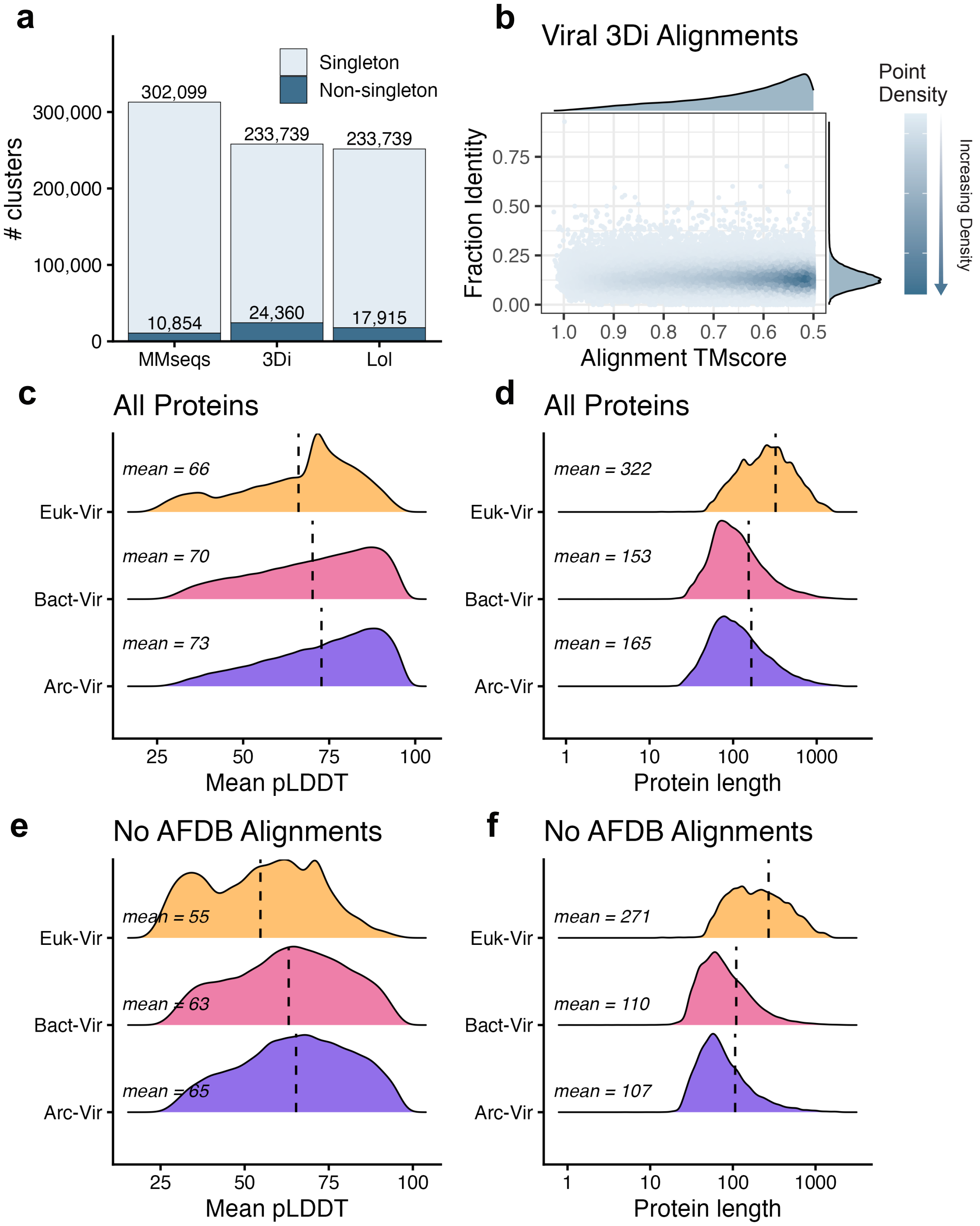

#### Extended Data Fig. 2

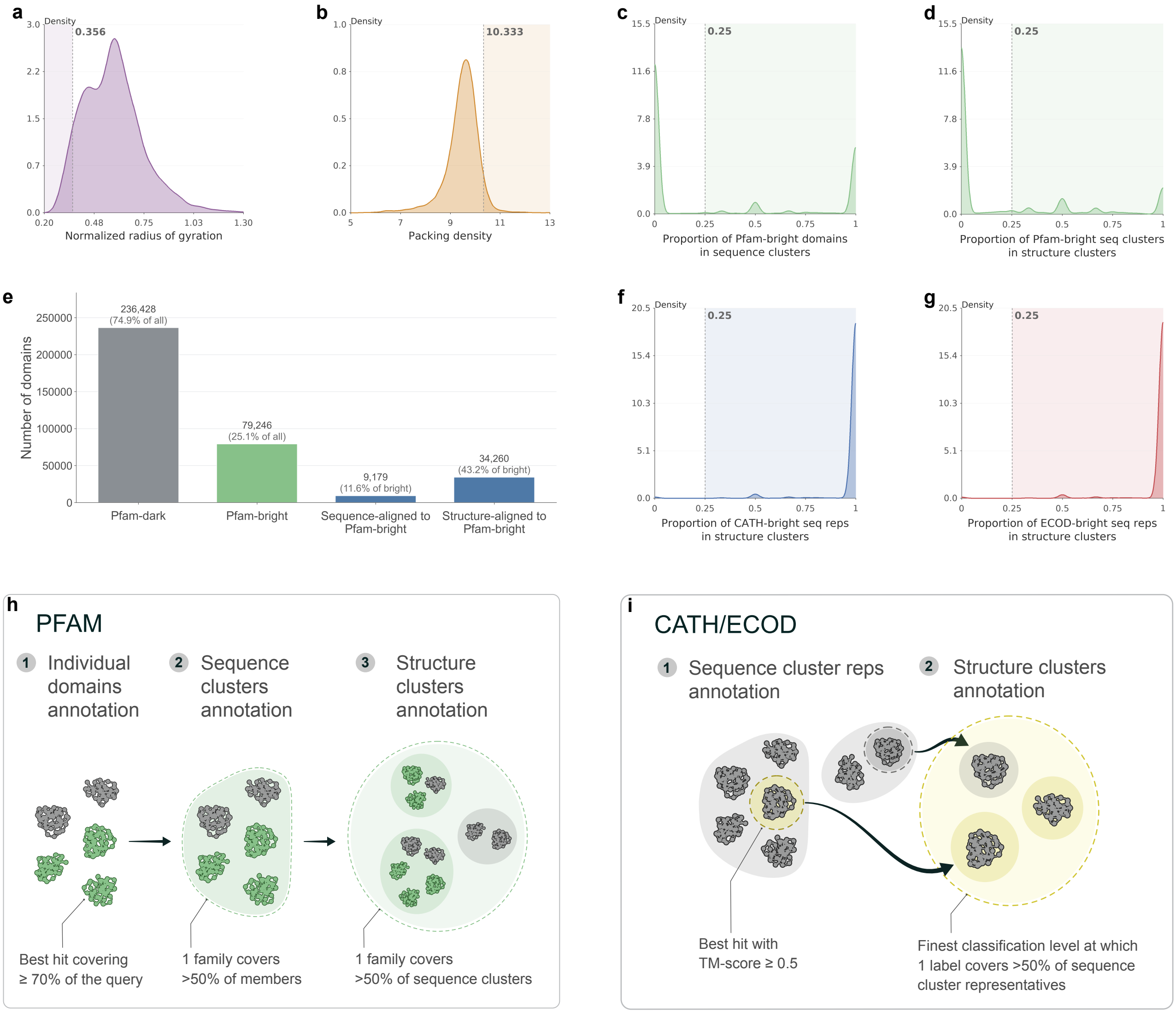

### Extended Data Fig. 3

**a**

**Mre11–Rad50-like fusion proteins**

| Identifier | Lineage | Host / organism | Confidence | Rad50 head evidence | Mre11 di-Mn ligands | Best Mre11 reference (TM-score) |
|---|---|---|---|---|---|---|
| **IMGVR_UViG_3300000153_000237…10016298**<br>IMGVR_UViG_3300000153_000237-3300000153-SI39nov09_135mDRAFT_10016298 | archaea | metagenomic | High | Zn hook | 6/8 | P. furiosus Mre11 (1II7)<br>TM 0.806 |
| **YP_009052356**<br>putative_DNA_repairing_ATPase__YP_009052356__Aureococcus_anophagefferens_virus__1474867 | NCLDV<br>Schizomimiviridae | Aureococcus anophagefferens virus | High | Walker-B | 7/8 | P. furiosus Mre11 (1II7)<br>TM 0.74 |
| **AID17555**<br>KJ591605.1__AID17555.1__X__00101 | phage<br>Caudoviricetes | Listeria phage LMTA-57 | High | Walker-B<br>+ Zn hook | 8/8 | P. furiosus Mre11 (1II7)<br>TM 0.731 |
| **YP_007006895**<br>NC_019521.1__YP_007006895.1__F405_gp016__00281 | phage<br>Caudoviricetes | Sphingomonas phage PAU | High | Zn hook | 7/8 | P. furiosus Mre11 (1II7)<br>TM 0.787 |
| **gwd2_scaffold_22…00196**<br>gwd2_scaffold_22_prodigal-single.1__X__X__00196 | phage | metagenomic | High | Zn hook | 7/8 | P. furiosus Mre11 (1II7)<br>TM 0.696 |
| **gwf1_scaffold_41…00459**<br>gwf1_scaffold_41_prodigal-single.1__X__X__00459 | phage | metagenomic | High | Zn hook | 7/8 | P. furiosus Mre11 (1II7)<br>TM 0.811 |
| **IMGVR_UViG_3300025676_000423…100663110**<br>IMGVR_UViG_3300025676_000423-3300025676-Ga0209657_100663110 | archaea | metagenomic | Medium | AAA_23 head | 6/8 | P. furiosus Mre11 (1II7)<br>TM 0.806 |
| **YP_007354557**<br>putative_DNA_repair_protein__YP_007354557__Acanthamoeba_polyphaga_moumouvirus__1269028 | NCLDV<br>Mimiviridae | Acanthamoeba polyphaga moumouvirus | Medium | AAA_23 head | 7/8 | P. furiosus Mre11 (1II7)<br>TM 0.78 |
| **YP_003987070**<br>uncharacterized_hydrolase__YP_003987070__Acanthamoeba_polyphaga_mimivirus__212035 | NCLDV<br>Mimiviridae | Acanthamoeba polyphaga mimivirus | Medium | AAA_23 head | 7/8 | P. furiosus Mre11 (1II7)<br>TM 0.747 |

**Intein variants of the complex**

| Identifier | Lineage / organism | Intein class | Block A (nucleophile) | Block G (His–Asn) | C-extein +1 | Splicing compatible | Hint TM | HEG (endonuclease) | Genome Mre11 |
|---|---|---|---|---|---|---|---|---|---|
| **PHAGE-A6--js4906-27-3_S30_scaffold_5…00192**<br>PHAGE-A6--js4906-27-3_S30_scaffold_5_curated_closed_complete_prodigal-single.1__X__X__00192 | phage<br>metagenomic | 1 | Cys | His–Asn | Ser | Yes | 0.817 | active<br>(TM 0.73) | — |
| **rifoxyc1_full_scaffold_3…00241**<br>rifoxyc1_full_scaffold_3_prodigal-single.1__X__X__00241 | phage<br>metagenomic | 1 | Cys | His–Asn | Ser | Yes | 0.939 | acidic subst.<br>(TM 0.827) | — |
| **IMGVR_UViG_3300025659_000107…10054961**<br>IMGVR_UViG_3300025659_000107-3300025659-Ga0209249_10054961 | archaea<br>metagenomic | 2 | not covered | His–Asn | Ser | Yes | 0.876 | none | — |
| **AIZ94713**<br>KP054477.1__AIZ94713.1__LfeInf_087__00087 | phage<br>Caudoviricetes<br>Lactobacillus phage LfeInf | 2 | Ala<br>(non-nucleoph.) | His–Asn | Ser | Yes | 0.813 | active<br>(TM 0.691) | intact |

**b**

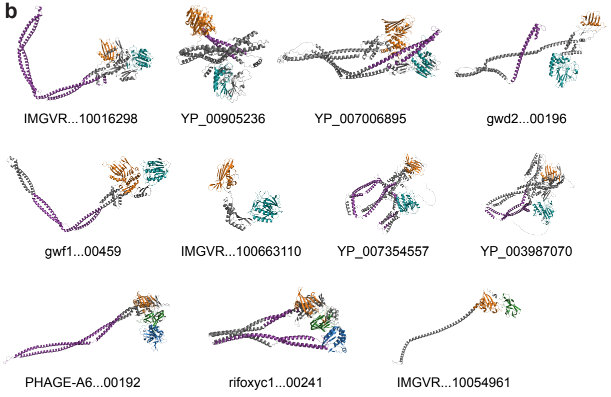


**c**

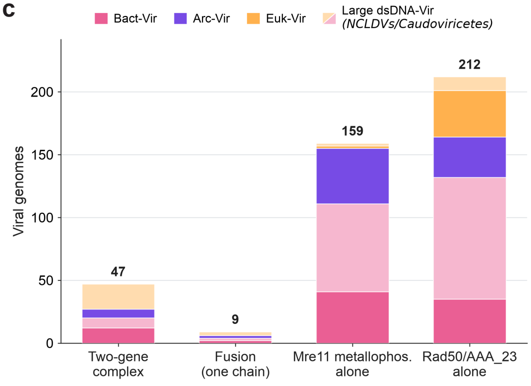

#### Extended Data Fig. 4

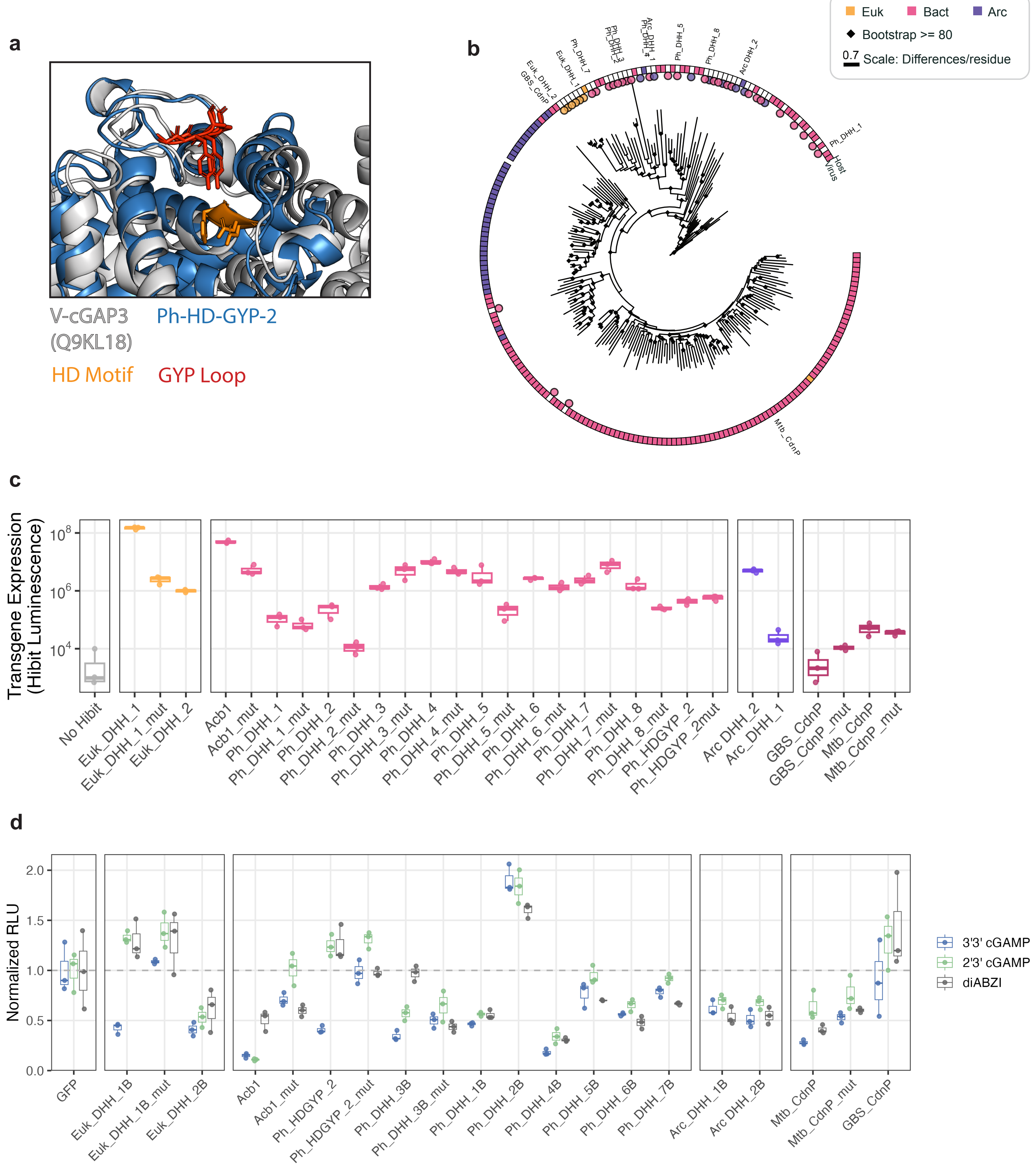

#### Extended Data Fig. 5

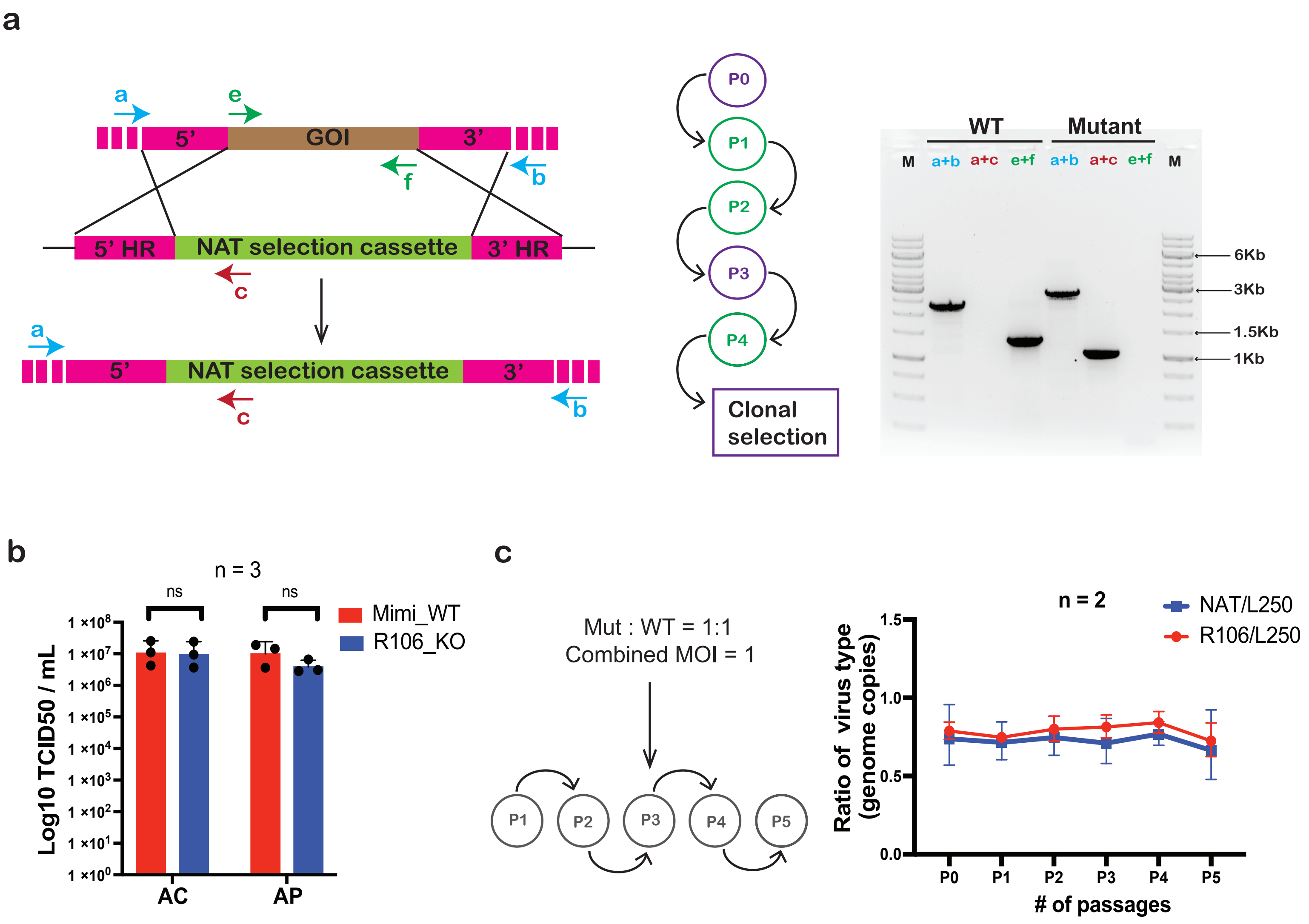

#### Extended Data Fig. 6

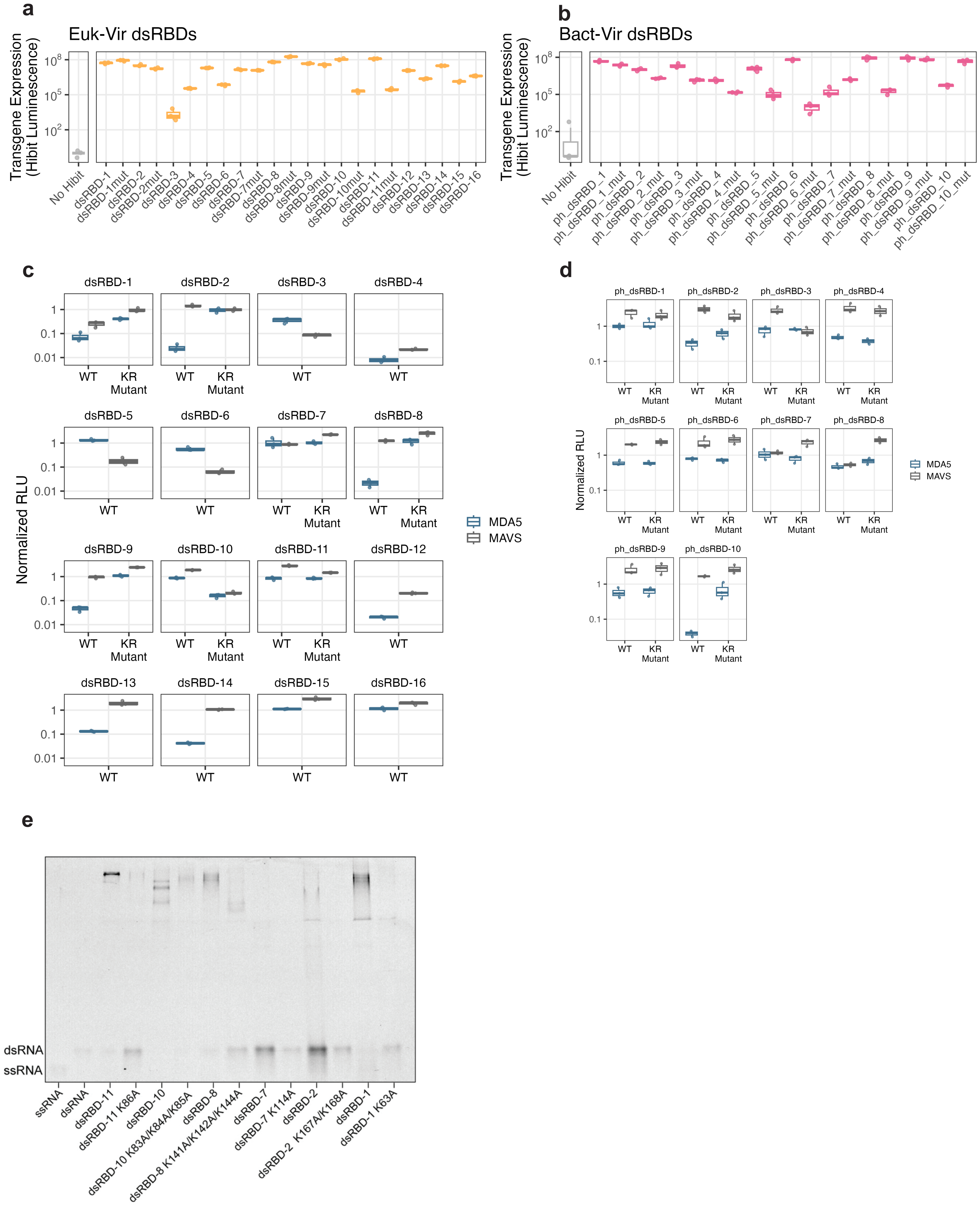
