## Supplementary figures for "Conserved folds enable immune antagonism across the tree of life"

**Supplementary Fig. 1.** Representative TLCs for substrate panel degradation analysis.

**Supplementary Fig. 2.** Binding analysis of dsRBDs versus KR mutant dsRBDs.

**Supplementary Fig. 3.** Source gel data.

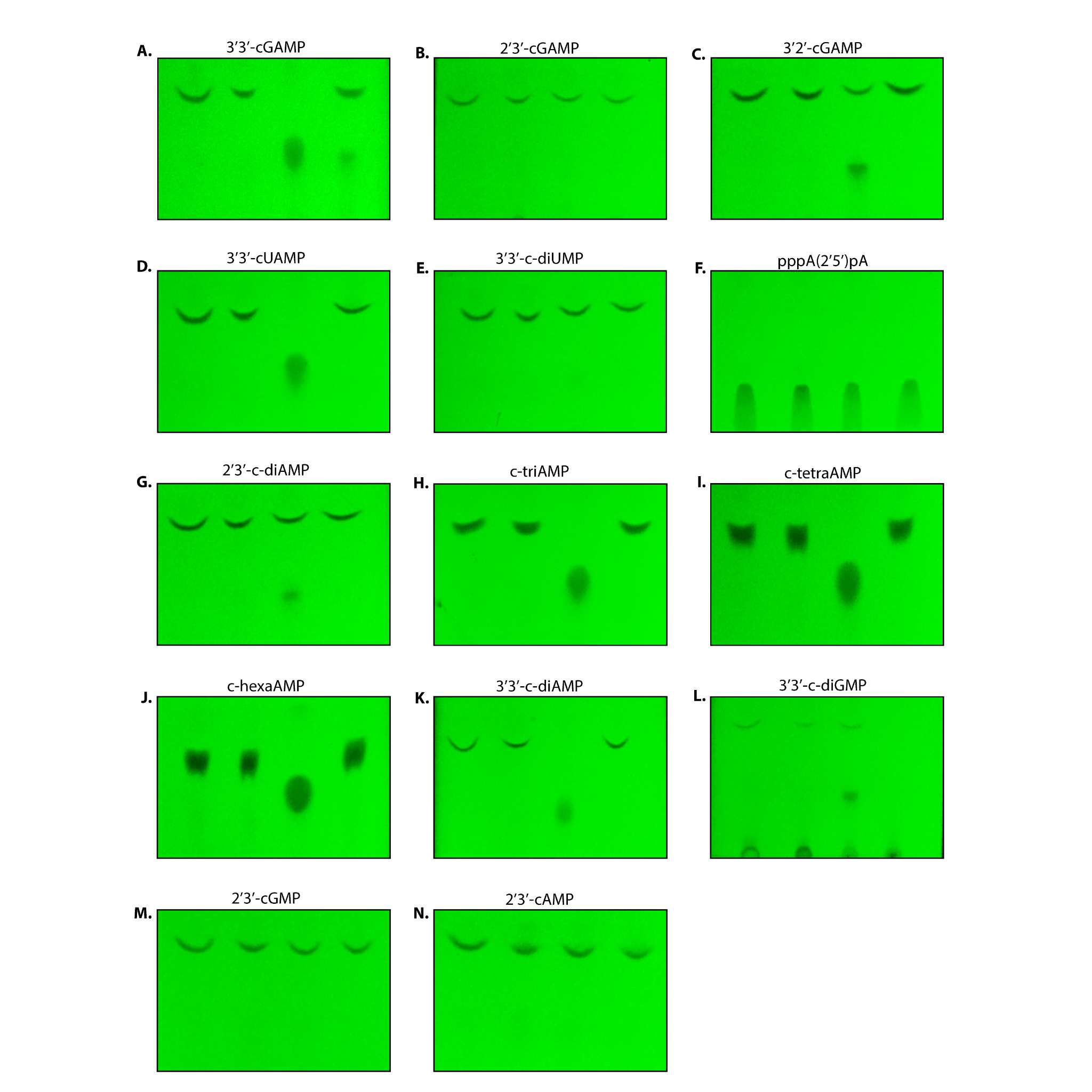
**Supplementary Fig. 1. Representative TLCs for substrate panel degradation analysis.** Each reaction was completed and assessed by TLC in technical duplicate. Lanes in order are no enzyme, a PDE candidate not reported in this work, Euk DHH_1B, and Ph HD-GYP_2, where: **A)** 3’3’-cGAMP **B)** 2’3’-cGAMP **C)** 3’2’-cGAMP **D)** 3’3’-cUAMP **E)** 3’3’-c-diUMP **F)** pppA(2’5’)pA **G)** 2’3’-c-diAMP **H)** c-triAMP **I)** c-tetraAMP **J)** c-hexaAMP **K)** 3’3’-c-diAMP **L)** 3’3’-c-diGMP **M)** 2’3’-cGMP **N)** 2’3’-cAMP.

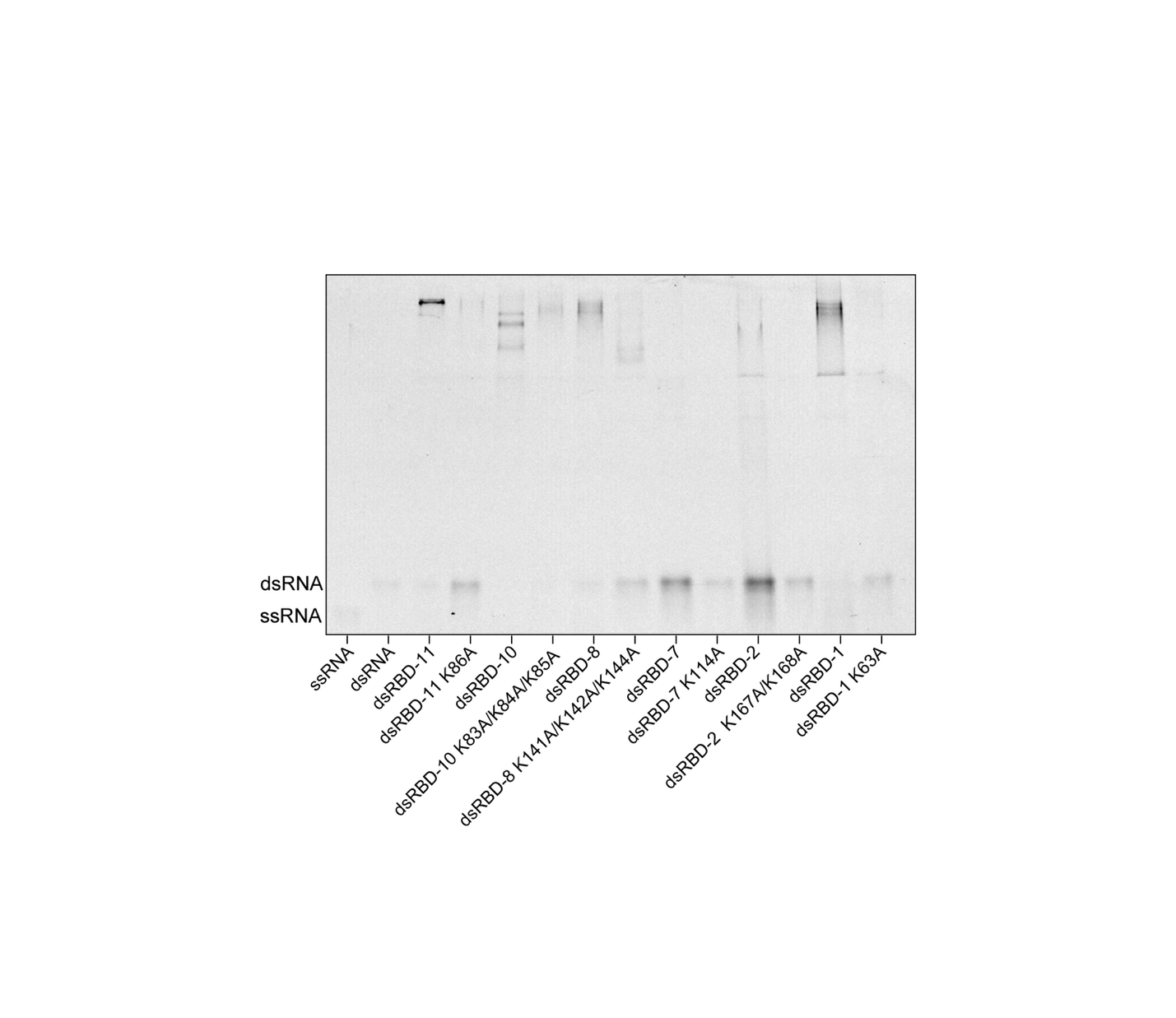

**Supplementary Fig. 3. Binding analysis of dsRBDs versus KR mutant dsRBDs.** Electrophoretic Mobility Shift Assay (EMSA; Methods) of candidate dsRBDs and K>A and/or R>A mutants.

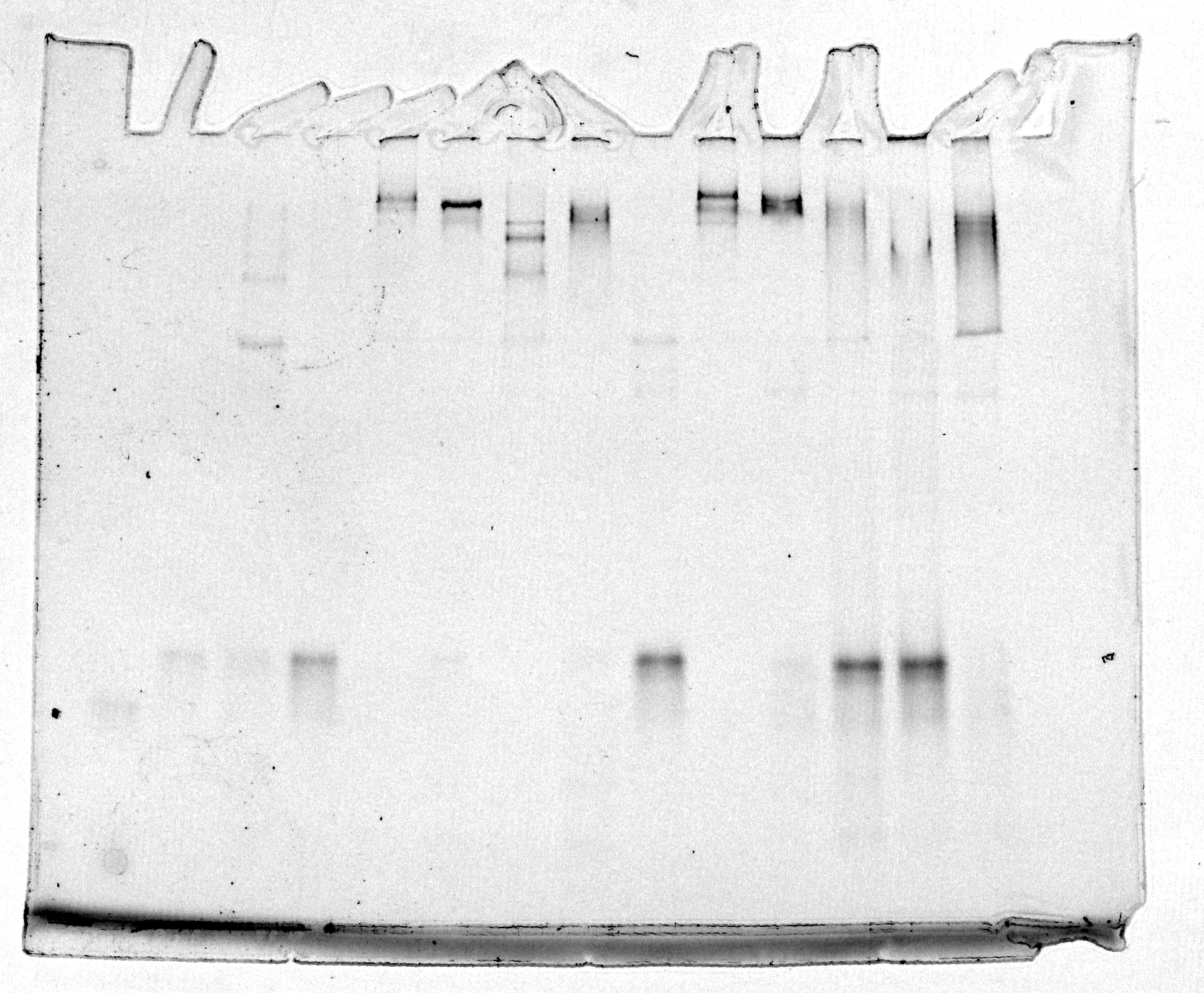

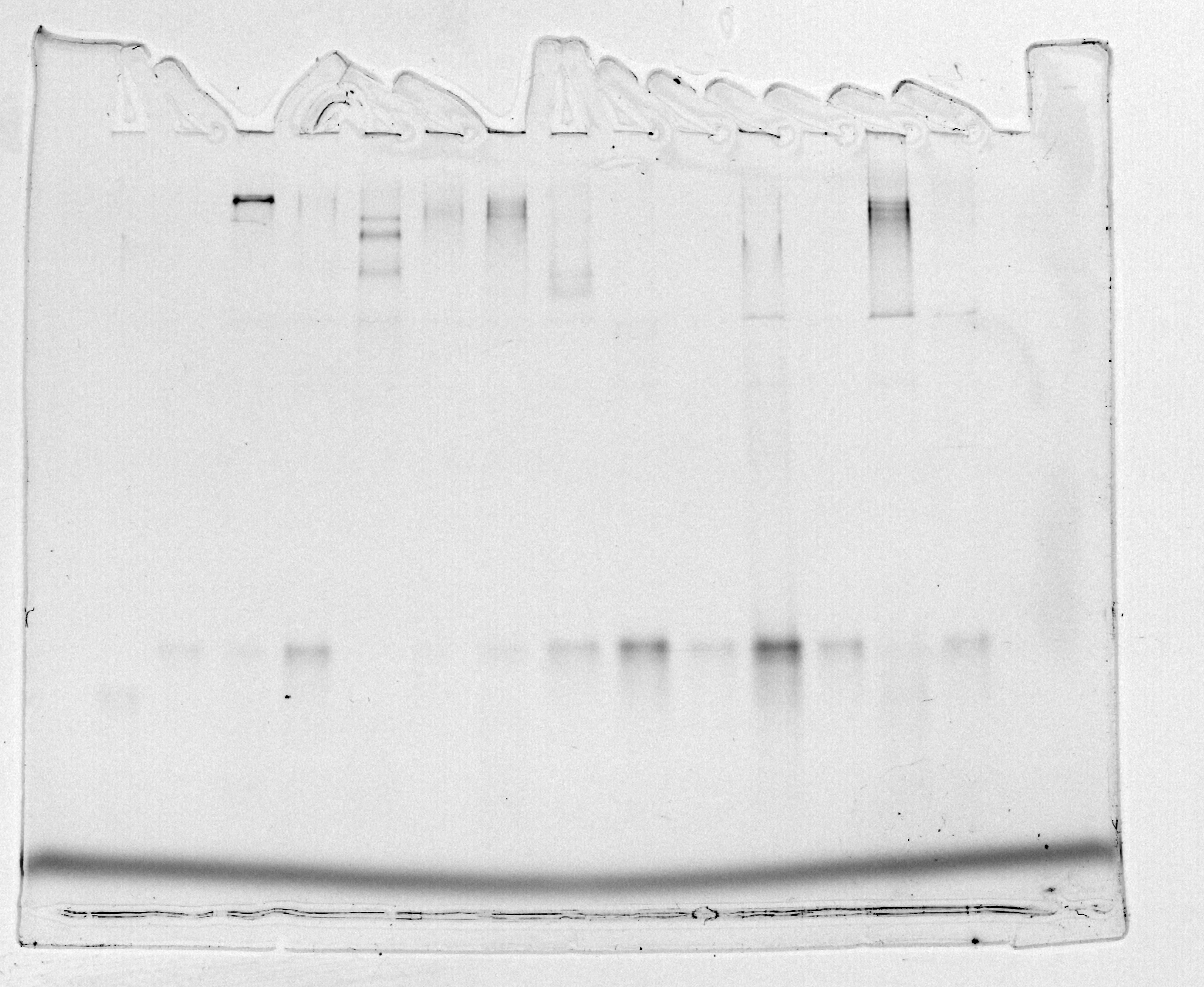

**Supplementary Fig. 4.** **Source gel data.**
